# CD47 is Required for Mesenchymal Progenitor Proliferation and Fracture Repair

**DOI:** 10.1101/2024.03.06.583756

**Authors:** Robert L. Zondervan, Christina A. Capobianco, Daniel C. Jenkins, John D. Reicha, Livia M. Fredrick, Charles Lam, Jeffery S. Isenberg, Jaimo Ahn, Ralph S. Marcucio, Kurt D. Hankenson

## Abstract

CD47 is a ubiquitous and pleiotropic cell-surface receptor. Disrupting CD47 enhances injury repair in various tissues but the role of CD47 has not been studied in bone injuries. In a murine closed-fracture model, CD47-null mice showed decreased callus bone volume, bone mineral content, and tissue mineral content as assessed by microcomputed tomography 10 days post-fracture, and increased fibrous volume as determined by histology. To understand the cellular basis for this phenotype, mesenchymal progenitors (MSC) were harvested from bone marrow. CD47-null MSC showed decreased large fibroblast colony formation (CFU-F), significantly less proliferation, and fewer cells in S-phase, although osteoblast differentiation was unaffected. However, consistent with prior research, CD47-null endothelial cells showed increased proliferation relative to WT cells. Similarly, in a murine ischemic fracture model, CD47-null mice showed reduced fracture callus bone volume and bone mineral content relative to WT. Consistent with our *in vitro* results, *in vivo* EdU labeling showed decreased cell proliferation in the callus of CD47-null mice, while staining for CD31 and endomucin demonstrated increased endothelial cell mass. Finally, WT mice administered a CD47 morpholino, which blocks CD47 protein production, showed a callus phenotype similar to that of non-ischemic and ischemic fractures in CD47-null mice, suggesting the phenotype was not due to developmental changes in the knockout mice. Thus, inhibition of CD47 during bone healing reduces both non-ischemic and ischemic fracture healing, in part, by decreasing MSC proliferation. Furthermore, the increase in endothelial cell proliferation and early blood vessel density caused by CD47 disruption is not sufficient to overcome MSC dysfunction.

## 1 Introduction

Cluster of differentiation protein 47 (CD47) is a 50 kDa protein that has five transmembrane regions with one extracellular end containing an IgV domain and an intracellular variably spliced C-terminal tail. ^1^ CD47 belongs to the immunoglobulin (Ig) superfamily and is expressed on the surface of essentially all human cells. ^2,3^

CD47 plays an important role in cancer biology, immunology, angiogenesis, and tissue regeneration. In particular, disruption of CD47 is beneficial in ischemic injuries to the liver^4–6^, kidney^7–10^, skin^11–14^, and brain. ^15^ Isenberg *et al*. demonstrated improved vascular remodeling, tissue viability, and perfusion after femoral artery ligation in the CD47-null mouse. ^12,13^ Furthermore, disruption of CD47 in endothelial cells (ECs) promotes proliferation via increased expression of stemness transcription factors (*cMyc*, *Klf4*, *Oct4*, and *Sox2*). ^16^

An important class of anti-angiogenic molecules are the secreted thrombospondins (TSPs) which are present in the fracture callus and bind to CD47. TSPs also bind to the extracellular matrix (ECM). ^17–19^ Unlike other proteins in the ECM, TSPs do not appear to serve a structural function, but rather modulate cell-matrix interactions via cell-surface receptors and cytokines. TSP1 and TSP2 are potent endogenous anti-angiogenic proteins that are over-expressed in non-healing wounds. ^20,21^ The mechanism of anti-angiogenesis of TSP1 and TSP2 is believed to be through interaction with cluster of differentiation 36 (CD36) and/or CD47.

In addition to its role in angiogenesis, CD47 has been implicated in the regulation of cellular differentiation. CD47 binds to cell surface signal regulatory protein alpha (SIRPα) and modulates mesenchymal progenitor cell (MSC) osteoblastogenesis. ^22^ Herein, we delineated the effects of an absence of CD47 on bone regeneration. Femoral fractures were studied in mice deficient in CD47 (CD47-null) and compared to wild-type mice (WT). *In vitro*, the impact of CD47 deficiency on marrow-derived and periosteum-derived cell proliferation and differentiation was investigated. The effect of CD47 disruption in ischemic tibial fractures was also assessed. Finally, to investigate transient disruption of CD47, WT mice with ischemic fracture were treated with CD47 morpholino oligonucleotide ([M]CD47) that temporarily limits mRNA translation. We report that loss of CD47 leads to a disruption of healing in both non-ischemic and ischemic fracture healing and that MSC lacking CD47 have reduced proliferation.

## 2 Results

### 2.1 Disruption of CD47 leads to delayed callus formation

We characterized femoral healing in CD47-null and WT mice at day 10 and 20 post-fracture using microcomputed tomography (µCT) as well as histomorphometric analysis. At day 10 post-fracture, there was evidence of decreased callus formation in CD47-null mice compared to WT (**Figure 1a**). Through uCT analysis, we observed reductions in bone volume and bone volume fraction, denoting a decrease in overall callus mineralization when CD47 was deleted (**Figure 1b-i**). Interestingly, bone formed in the absence of CD47 showed a higher average density (TMD) at day 10 (**Figure 1i**). Through histomorphometry, we further observed a decrease in callus parameters including the percentage of bone and marrow when CD47 was deleted, but an increase in fibrous tissue (**Figure 2a-i)**. This corroborates the uCT data of a reduction or delay in fracture callus formation in the absence of CD47. At day 20 post-fracture (**Figure 1 & 2)**, we noted that these reductions in callus size are no longer apparent. This led us to suspect that the early cell signaling responses during fracture healing are perturbed in the absence of CD47.

**Figure 1.**
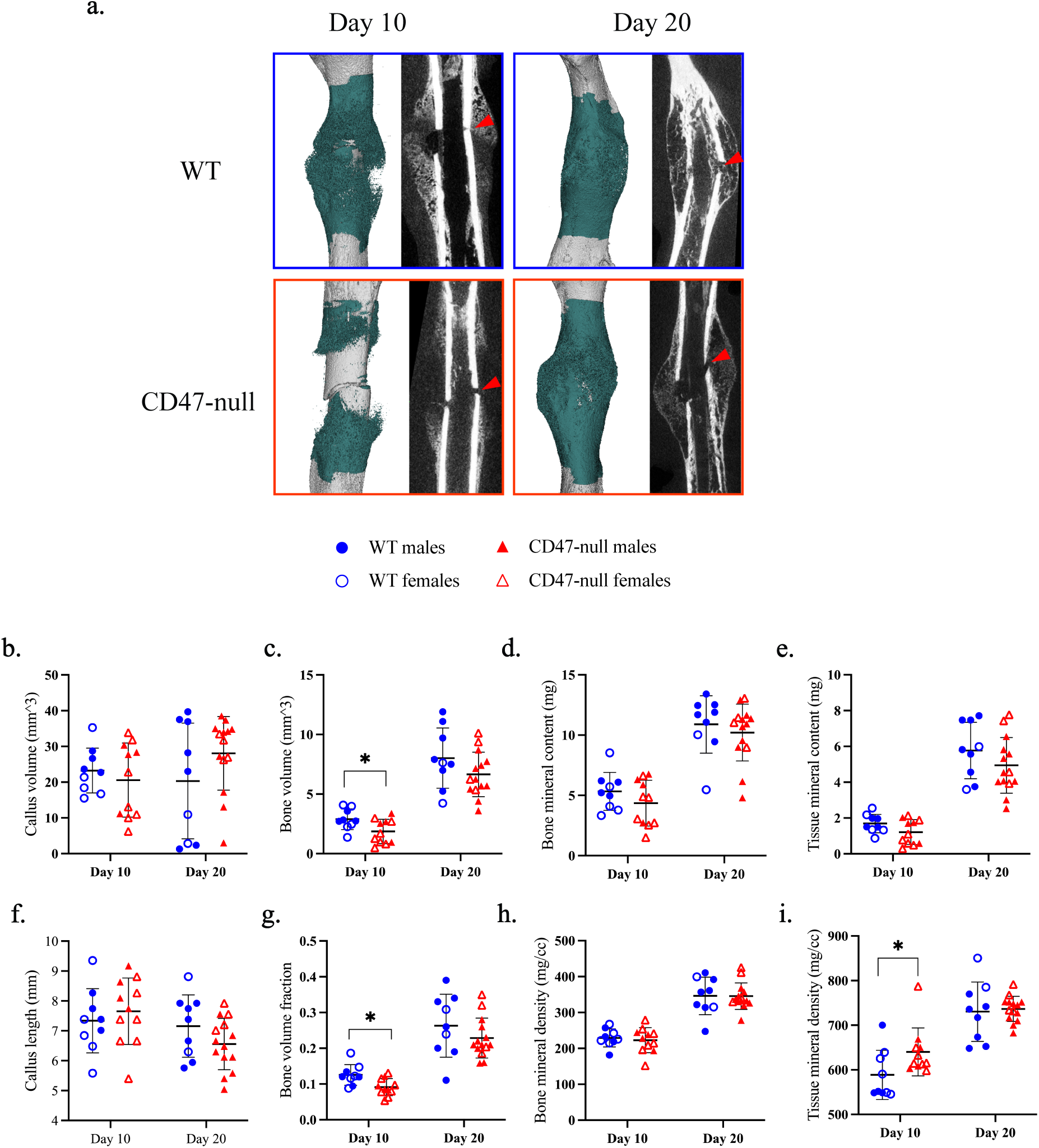
CD47-null fractures show reduced bone formation, but increased tissue mineral density. µCT analysis of transverse femoral fracture in WT (n=9) and CD47-null (n=11-14) mice at 10- and 20-days post-fracture. **a**, Representative 3D reconstructions (white background) and sagittal-plane reconstruction (black background) of WT and CD47-null mice at days 10 and 20 post-fracture. Location of fracture (red arrowhead) is marked on the sagittal reconstructions. Representative 3D reconstructions include callus mineralization (teal). **b-i,** Callus morphology (mean±SD) at day 10 and 20 post-fracture. \**P<*0.05, two-sided t test performed at each timepoint.

**Figure 2.**
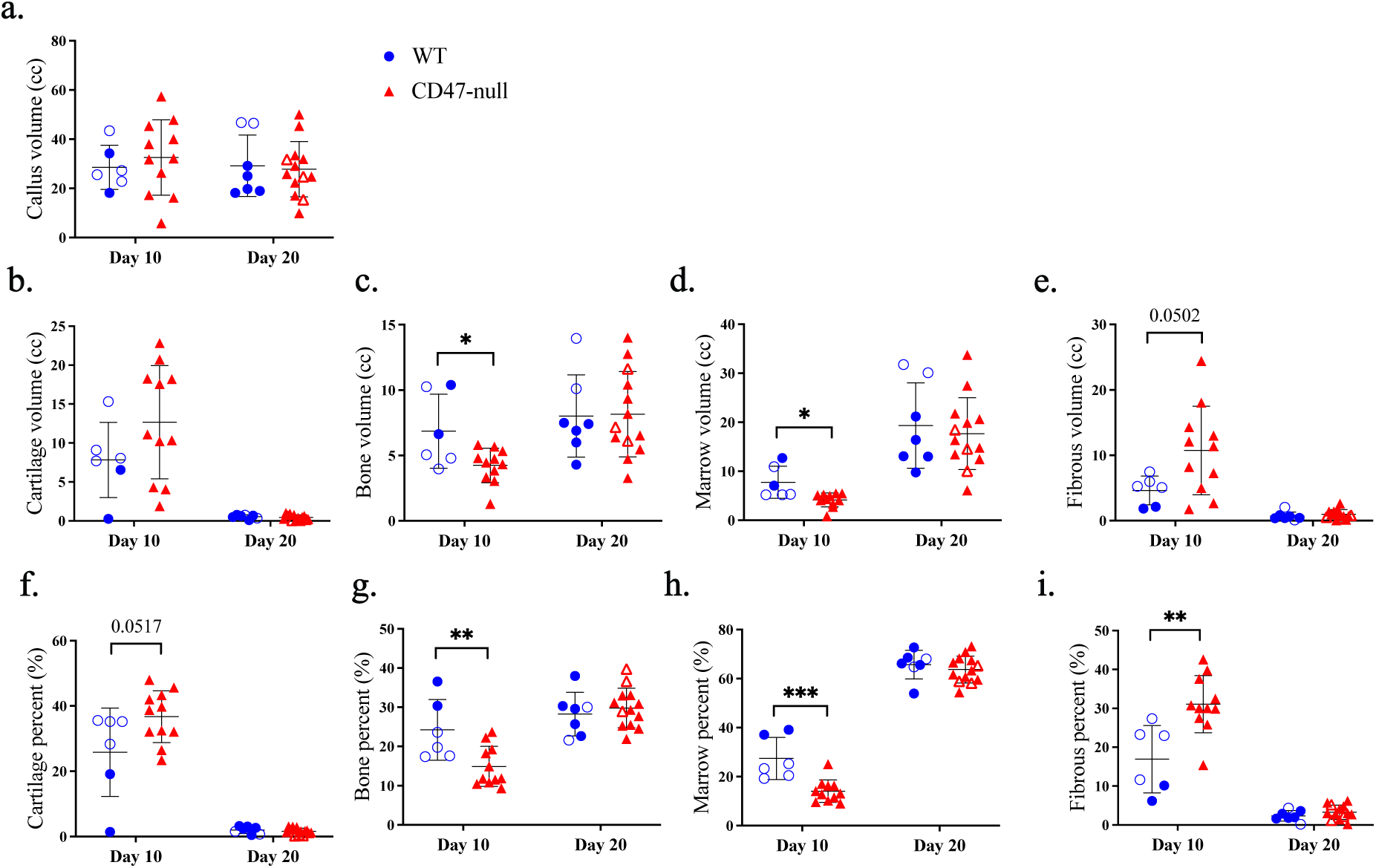
Genetic knockout of the thrombospondin-CD47 axis has minimal effect on late fracture callus composition. Histomorphometry of transverse femoral fractures in WT (n=6-7), and CD47-null (n=11-13) mice at 10- and 20-days post-fracture. **a**-**i**, Callus composition quantified through histomorphometry (mean±SD) at day 10 and 20 post-fracture. \**P<*0.05, \*\**P<*0.01, \*\*\**P<*0.001, two-sided t test performed at each timepoint.

### 2.2 Marrow-derived mesenchymal progenitor cell colony expansion and proliferation is reduced in CD47-null mice

Based on the phenotype of less bone forming during early fracture healing in CD47-null mice, we next studied the cells that contribute to callus maturation. Activation and rapid expansion of mesenchymal progenitor cells (MSC) from the marrow and periosteum is necessary for proper callus formation. After 12 days in culture, both WT and CD47-null marrow-derived MSC formed colonies (**Figure 3a**). However, there was a difference in colony phenotype (**Figure 3b**). WT and CD47-null cells produced an equivalent number of small (25-100 ALP positive cell) colonies, but WT produced significantly more colonies that were judged large (>100 ALP positive cell). This suggested a role for CD47 regulating proliferative capacity or contact inhibition in the absence of CD47. Proliferation was investigated indirectly using the MTT assay to measure cellular metabolism. WT cells had significantly higher proliferation at days 6 and 10 post-harvest (**Figure 3c**). There was no difference immediately after plating and at 14-days post-harvest which suggests CD47 promotes MSC proliferation within the conditions of the experiment. To further investigate the proliferative phenotype of CD47-null MSC, cells were passaged to select for adherent cells (MSC) over non-adherent cells present in whole-marrow aspirate. MTT as a surrogate for proliferation was measured at day 3, 6, and 9 in post-passage. There was a significant increase in proliferation in WT over CD47-null MSC at all time points (**Figure 3d**). Results were confirmed using BrdU incorporation which was measured on days 1 and 12 post-passage. BrdU incorporation was lower in CD47-null cells at day 1 (t=4.469, p=0.0012, two-sided t test) and day 12 (t=6.872, p<0.0001, two-sided t test) post-passage (data not shown).

**Figure 3.**
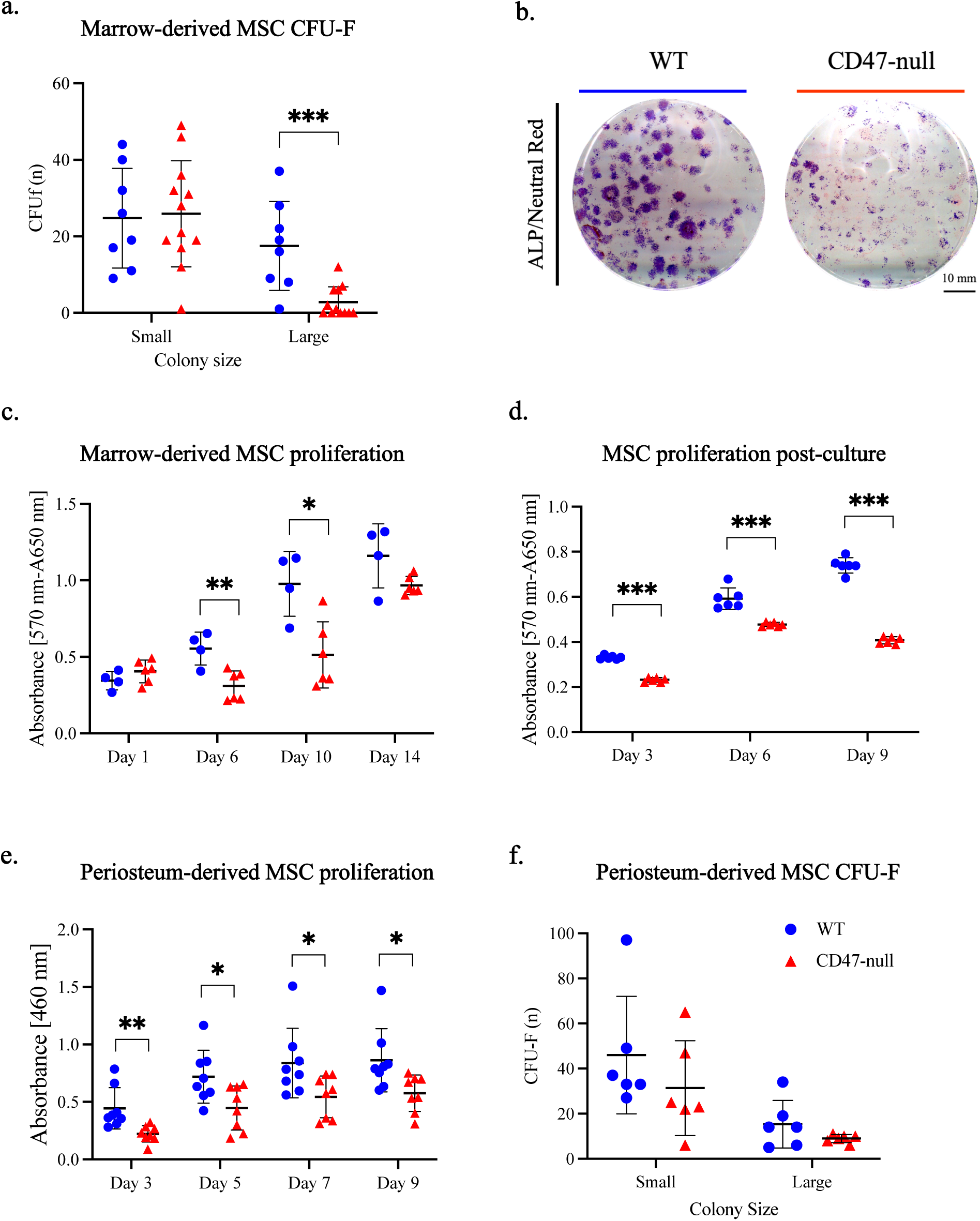
CD47 is required for cell colony expansion and proliferation. Marrow and periosteal MSC harvested from the femur and tibia of WT and CD47-null mice. **a,** CFU-F of WT (n = 8) and CD47-null (n = 12) marrow-derived cells. **b,** Representative CFU-f plate from WT (left) and CD47-null (right) mice. **c**, MTT assay of marrow-derived cells MSC from WT (n= 4) and CD47-null (n=4) mice at 1, 6, 10, and 14 days in culture. **d**, MTT assay of marrow MSC from WT (n= 6) and CD47-null (n=6) after 1 passage at 3, 6, and 9 days in culture. **e,** CCK8 assay of periosteal MSC from WT (n= 8) and CD47-null (n=8) after 1 passage at 3, 5, 7, and 9 days in culture. **f,** CFU-F of periosteal MSC from WT (n= 6) and CD47-null (n=6) mice. Mean ±SD, \**P<*0.05, \*\**P<*0.01, \*\*\**P<*0.001 two-sided t tests.

### 2.3 Periosteal-derived MSC colony expansion and proliferation is reduced in CD47-null mice

To investigate the characteristics of periosteum-derived MSC in comparison to the marrow-derived MSC, CFU-F expansion and CCK8 proliferation assays were performed. Proliferation was assessed at 3, 5, 7, and 9 days in periosteal MSC after plating. There was a significant decrease in CD47-null periosteal MSC proliferation at all timepoints (**Figure 3e)**. CFU-F analysis of alkaline phosphatase positive colonies in periosteal MSC revealed a trend of reduced colony formation, particularly of larger ALP positive colonies within CD47-null cells (**Figure 3f)**. Results in marrow-derived and periosteum-derived MSC indicate that CD47 supports proliferation but is not required for initial colony formation.

### 2.4 CD47-null MSC show reduced cell cycle progression

To explore mechanism that may affect proliferation of CD47-null MSC, cultured cells were assayed for disruptions in gene expression, apoptosis, and cell cycle regulation. Gene expression analysis confirmed complete disruption of CD47 expression in CD47-null cells and a decrease in *Oct4*, *Klf4*, and *cMyc* (markers for stem cells), and alkaline phosphatase (*ALP*) (a marker for pre-osteoblasts) (**Supplemental Figure 1a-e**). CD47-null cells also expressed decreased levels of Caspase3, suggesting the decrease in proliferation is not due to apoptosis (**Supplemental Figure 1f**). Similarly, assessment of apoptosis by flow cytometry showed that there were no differences in either apoptosis or necrosis between WT and CD47-null cells (**Supplemental Figure 2**). CD47-null cells also demonstrated decreased levels of *FOXm1*, a factor required for proliferation and differentiation (**Supplemental Figure 1g**) ^23^. Cell cycle analysis by flow cytometry revealed a 2.7-fold decrease in percent of CD47-null MSC in S phase (**Figure 4**).

**Figure 4.**
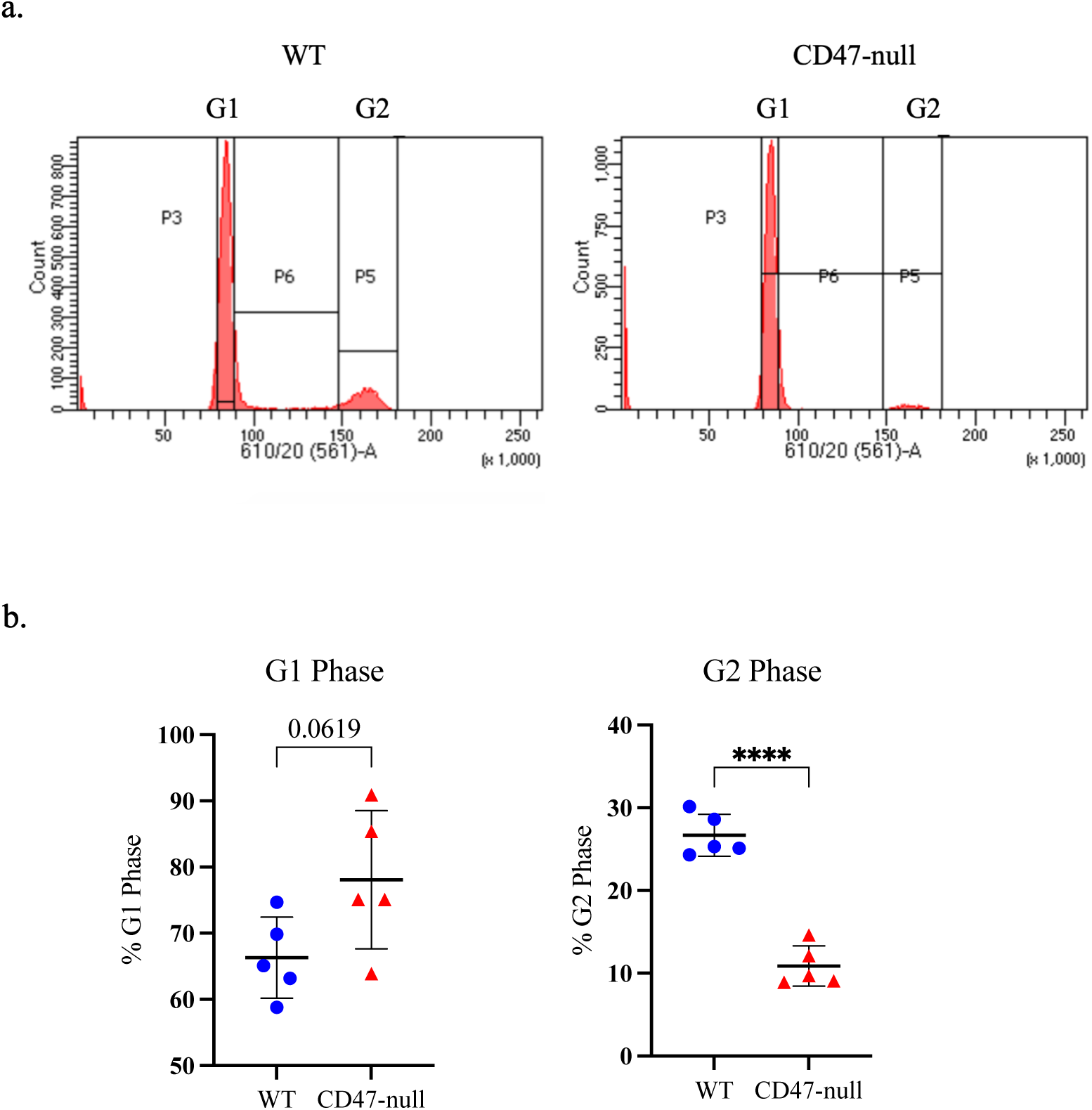
Loss of CD47 decreases percent of MSC in S/G2 phase. Cell cycle analysis of marrow-derived MSC harvested from the femur and tibia of WT (n=5) and D47-null (n=5) mice. **a,** Representative histograms of WT (left) and CD47-null (right) with cell cycle analysis overlays **b**, Percent of cells in G1 phase or S/G2. Mean ±SD*, *P<*0.05, \*\**P*<0.01, \*\*\**P*<0.001, two-sided t test.

### 2.5 Mesenchymal stem cell osteoblastic differentiation in CD47-null cells

To assess the ability of CD47-null cells to differentiate into mineral producing osteoblasts, marrow cells were harvested from the femur and tibia of WT and CD47-null mice. After 7 days, cells were passaged into osteopermissive media. At day 14, plates were stained with either Alizarin red S to observe mineralization or ALP/Neutral Red. ALP is indicative of osteoblast differentiation while Neutral Red will stain all cells, regardless of ALP positivity (**Supplemental Figure 3**). Disruption of CD47 decreased the overall number of cells, as previously shown, and concomitantly, the amount of mineralization was significantly lower in CD47-null cells (**Supplemental Figure 3**). Interestingly, when normalized by cell density using ALP (as most cells are ALP positive), there was no difference between WT and CD47-null mineralization, suggesting that an absence of CD47 does not negatively impact osteoblast differentiation, but rather decreases the number of cells that are differentiating to become osteoblasts.

### 2.6 The basal distribution of periosteal skeletal stem cells is not different between WT and CD47-null mice

To examine the skeletal stem cell niche^24^, we performed flow cytometry to analyze skeletal stem cell populations present within the periosteum of WT and CD47-null mice. Flow cytometry confirmed the majority of cells within the periosteum are CD51+. Distribution of periosteal cell populations across mouse skeletal stem cells (mSSC) (CD51+/Ly51-/CD90-/CD105-/CD200+), pre-bone cartilage stromal progenitors (pBCSP) (CD51+/Ly51-/CD90-/CD105-/CD200-), and bone cartilage stromal progenitors (BCSP) (CD51+/Ly51-/CD90-/CD105+) revealed around 40% of cells within the periosteum to be of pre-bone cartilage progenitor immunotype along with approximately 20% of cells being stem cells. There were no appreciable differences in total cell numbers of cells in the niche of uninjured bone and similarly, percentage distributions were not different between CD47-null and WT **(Supplemental Figure 4).**

### 2.7 CD47-null endothelial cells show increased proliferation

TSP1-CD47 signaling leads to cellular senescence in endothelial cells, and in the absence of TSP1 and CD47, endothelial cells show increased proliferation^25^, as opposed to our observation of decreased proliferation of MSC. Thus, we wanted to establish in our laboratory the proliferative capacity of CD47-null endothelial cells. Lungs were harvested from 4-week-old mice and digested in collagenase A before sorting for CD31+ cells using magnetic beads for cell isolation (**Figure 5a**). Purity of the isolate was confirmed using flow cytometry (**Figure 5b**). Prior to sorting, 11.65% of harvested lung cells were positive for CD31. After CD31 MicroBead sorting, the percent of CD31+ isolated cells increased to 96.44%. CD31+ cells were plated on gelatin coated 12-well plates at a density of 100,000 cells per well. Cellular proliferation of WT and CD47-null endothelial cells was determined at day 3, 6, and 9 with an MTT assay. The relative proliferation of CD47-null endothelial cells was 46.5%, 39.5%, and 32.8% higher than WT cells at days 3, 6, and 9, respectively (**Figure 5c**). However, only the difference in absorbance at day 3 was significant. Still, there was a significant overall effect of genotype and day suggesting loss of CD47 increased proliferation of endothelial cells, consistent with the well-established phenotype. Of note, in this model system, apoptosis of the endothelial cells results in decreases in MTT over time in culture, irrespective of genotype (**Figure 5c**).

**Figure 5.**
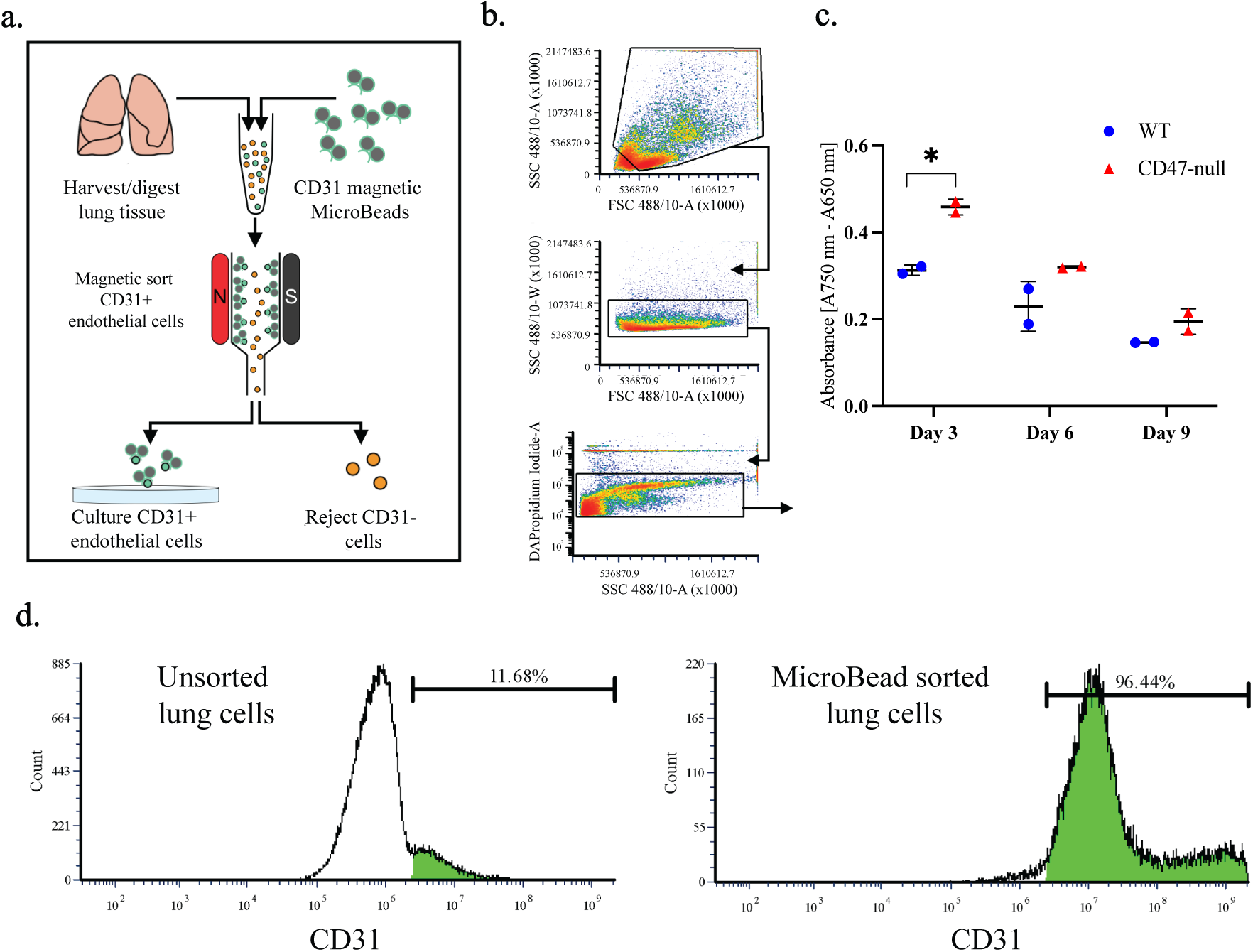
Loss of CD47 promotes endothelial cell proliferation. **a**, Schematic of endothelial cell isolation and culture from the lungs of 4-week-old mice. **b**, Flow cytometry gating scheme for lung cell density (top), double discrimination (middle), and alive/dead (bottom). **c,** MTT assay of WT and CD47-null endothelial cells after 3, 6, and 9 days in culture. **d**, Representative histograms of CD31+ lung cells before CD31 MicroBead sorting (left) and after (right). Mean ±SD, \**P<*0.05, two-sided t test.

### 2.8 Baseline vascular function and response to ischemic fracture in CD47-null mice is similar to WT mice

Prior work has shown that disruption of CD47 increases vascularization in non-bone tissues and increases healing of ischemic injuries. We sought to investigate the effect of disrupting CD47 in ischemic fracture healing. Prior to the ischemic fracture, WT and CD47-null mice showed equivalent levels of hind limb perfusion (**Supplemental Figure 5a**). Immediately following ischemic surgery and fracture the right (nonischemic) to left (ischemic) limbs were compared which demonstrated a significant mean decrease of 141.3±7.14 and 129.1±8.23 PU in WT and CD47-null mice, respectively. The difference in non-ischemic limb perfusion after surgery is likely a byproduct of being under anesthesia for ∼30 minutes. To compare between WT and CD47-null animals, perfusion was normalized to control for intra- and inter-animal differences. A relative perfusion change was generated by calculating the percent change in ratio of the ischemic limb to the non-ischemic limb. Immediately following surgery there was no significant difference in relative perfusion change between WT and CD47-null mice (**Supplemental Figure 5b**). Ischemic surgery induced a relative decrease of 81.83±1.03 and 82.04±1.45 in WT and CD47-null mice, respectively. These results suggest there is no baseline functional vascularity difference in CD47-null mice and no difference in immediate recovery or response.

### 2.9 Disruption of CD47 in early ischemic fracture impairs callus formation

After ischemic surgery and tibial fracture, fractured limbs were harvested at day 10, 15, and 20 post-fracture to characterize the callus phenotype at various stages of callus formation. At day 10 post-fracture, there was minimal development of a callus. Early signs of callus formation appeared as thickening of the periosteum distant to the fracture site both proximally and distally. There was no significant difference in callus composition at day 10 (**Figure 6a-i**), but CD47-null mice began to show lower bone mineral content (**Figure 6d**). At day 15 post-fracture, disruption of CD47 began to significantly impair callus formation. CD47-null mice had a total callus volume 47.8% smaller than WT mice. Compared to WT mice (**Figure 6b**), CD47-null mice also had reduced bone volume (-39.8%) (**Figure 6c**), bone mineral content (-47.2%) (**Figure 6d**), and tissue mineral content (-43.0%) ((**Figure 6e**). However, despite measured differences in absolute volumes and amounts, when normalized by callus volume (bone volume fraction) (**Figure 6g**), there was no difference between WT and CD47-null mice. The bone mineral density and tissue mineral density showed no difference between WT mice (**Figure 6f-i**). The differences observed in early callus formation between WT and CD47-null mice resolve by day 20 post-fracture. No defined differences were observed through histomorphometry at day 10 and 15 despite these changes noted via µCT (**Supplemental Figure 6).** This likely reflects technical differences between µCT and histology, as µCT can detect mineralized voxels (bone) at high resolution, but cannot as easily determine callus size, and similarly, histological assessment of bone is less sensitive than µCT.

**Figure 6.**
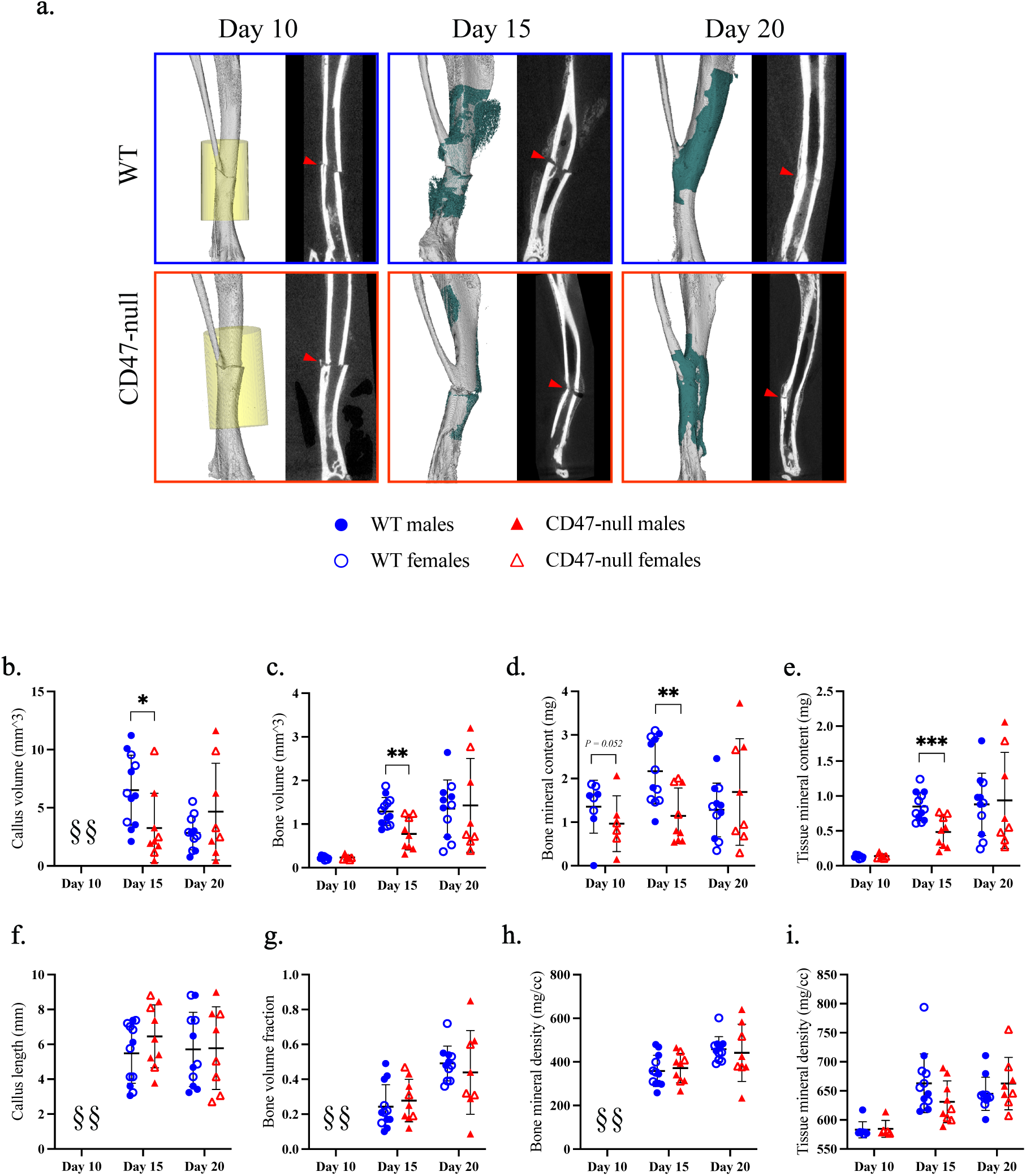
Genetic knockout of CD47 inhibits early ischemic fracture callus formation. µCT analysis of ischemic tibia fracture callus of WT (n=7-11) and CD47-null (n=6-9) mice at day 10, 15, and 20 post-fracture. **a**, Representative 3D reconstructions (white background) and sagittal-plane reconstruction (black background) of WT (top row) and CD47-null (bottom row) mice at day 10, 15, and 20 post-fracture. Location of fracture (red arrowhead) is marked on the sagittal reconstructions. Day 10 3D reconstruction includes representative cylindrical ROIs (transparent yellow cylinder) used to calculate callus morphology. Day 15 and 20 representative 3D reconstruction include highlighted callus mineralization (teal). **b-i**, Callus morphology (mean±SD) at days 10, 15, and 20 post-fracture. \**P<*0.05, \*\**P<*0.01, \*\*\**P<*0.001, two-sided t tests performed at each timepoint; § no data.

### 2.10 CD47-null mice do not show enhanced perfusion following ischemic injury

Next, we considered how WT and CD47-null mice recovered from an ischemic injury and if the callus phenotype was a result of prolonged ischemia or cellular dysfunction. Another group of WT and CD47-null mice were used for this experiment. After the initial ischemic fracture, serial measurements of perfusion were performed daily using laser doppler flowmetry of the plantar surface of the hindfoot. Using experimental perfusion results from day 0 through day 9 post-ischemic injury, the data was fit to a one-phase nonlinear equation:

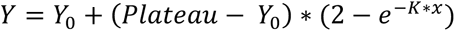

where *Y* is relative perfusion change, *Plateau* is the offset, *K* is the rate of perfusion change, and *x* is time. The resulting fits for WT (*R*^2^=0.7609) and CD47-null (*R*^2^=0.6223) were:

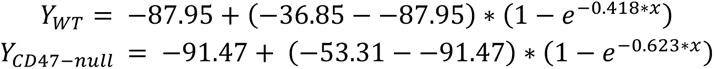

suggesting CD47-null mice recover faster, but maximum recovery is limited to 36.85% of baseline perfusion compared to WT which can recover to 53.32% of baseline perfusion (*F*_3,74_=19.87, *p<*0.0001, extra sum-of-squares F test) (**Figure 7a)**.

**Figure 7.**
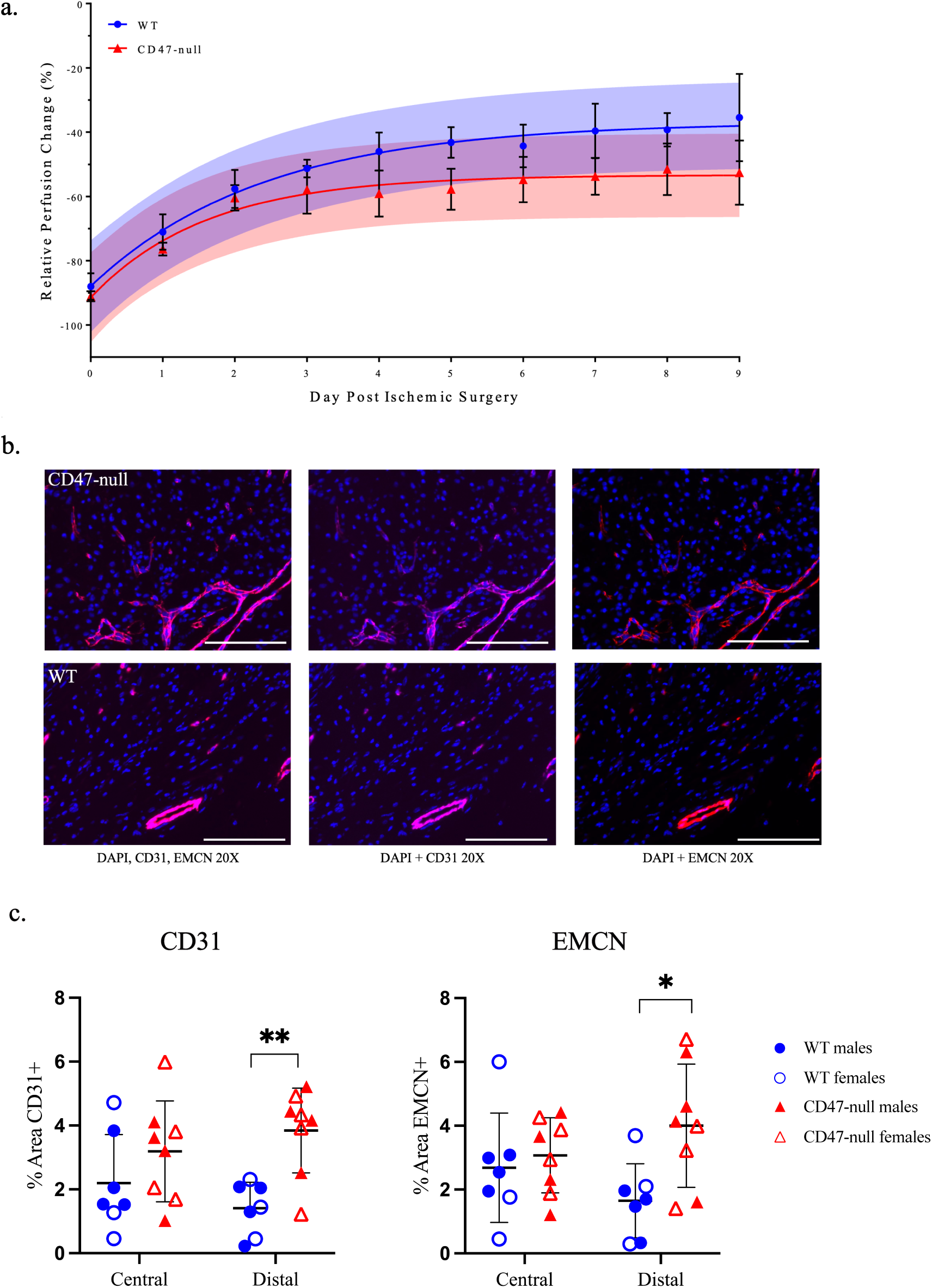
Genetic knockout of CD47 limits recovery of whole limb perfusion, but shows local increases in endothelial cells after induced ischemia. Relative perfusion WT and CD47-null mice at days 0-9 post-ischemic surgery using laser doppler flowmetry. **a,** Perfusion data was fit to a one-phase non-linear curve for WT (fit is the solid blue line with 95% CI in transparent blue; *R^2^*=0.7609) and CD47-null (fit is the solid red line with 95% CI in transparent red; *R^2^*=0.6223). The rate of recovery was faster, but maximum recovery was significantly lower in CD47-null compared to WT mice (P<0.0001, extra sum-of-squares F test). **b,** Representative immunofluorescence staining of CD31 and EMCN in CD47-null (n=7) and WT (n=8) ischemic fractures at 20X (Scale bar = 100 µm). **c,** CD31 quantification in WT and CD47-null ischemic fracture calluses at day 4 post-fracture at peripheral and central regions of the fracture callus. **D,** EMCN quantification in WT and CD47-null ischemic fracture calluses at day 4 post-fracture at peripheral and central regions. Mean ±SD, \**P<*0.05, \*\**P<*0.01, two-sided t test.

### 2.11 Endothelial cell density increases in the absence of CD47

Callus endothelial cell density was calculated at day 4 post-fracture. The callus was divided into two regions, central and distal relative to the fracture, for analysis. In CD47-null mice there is higher density of EMCN+ and CD31+ cells in the peripheral callus, while the central callus region did not reflect this difference **(Figure 7b-c)**. This data shows that disruption of CD47 leads to an increase in endothelial cell density at the periphery of the callus, consistent with our evaluation of isolated endothelial cells *in vitro*. We qualitatively observed a strong overlap between EMCN+ and CD31+ cells, which are termed H vessels. ^26^

### 2.12 CD47 disruption correlates with decreased cell proliferation

To investigate cell proliferation in the early ischemic fracture callus, WT and CD47-null mice were injured and harvested at day 4 and 7. Mice were EdU labeled prior to harvest. Immunofluorescence revealed no difference in EdU incorporation between the WT and CD47-null ischemic fracture at day 4 (**Figure 8a & 8b**), however there is a decrease in EdU+ cells within the ischemic fracture callus at day 7 under CD47-null conditions **(Figure 8c & 8d)**.

**Figure 8.**
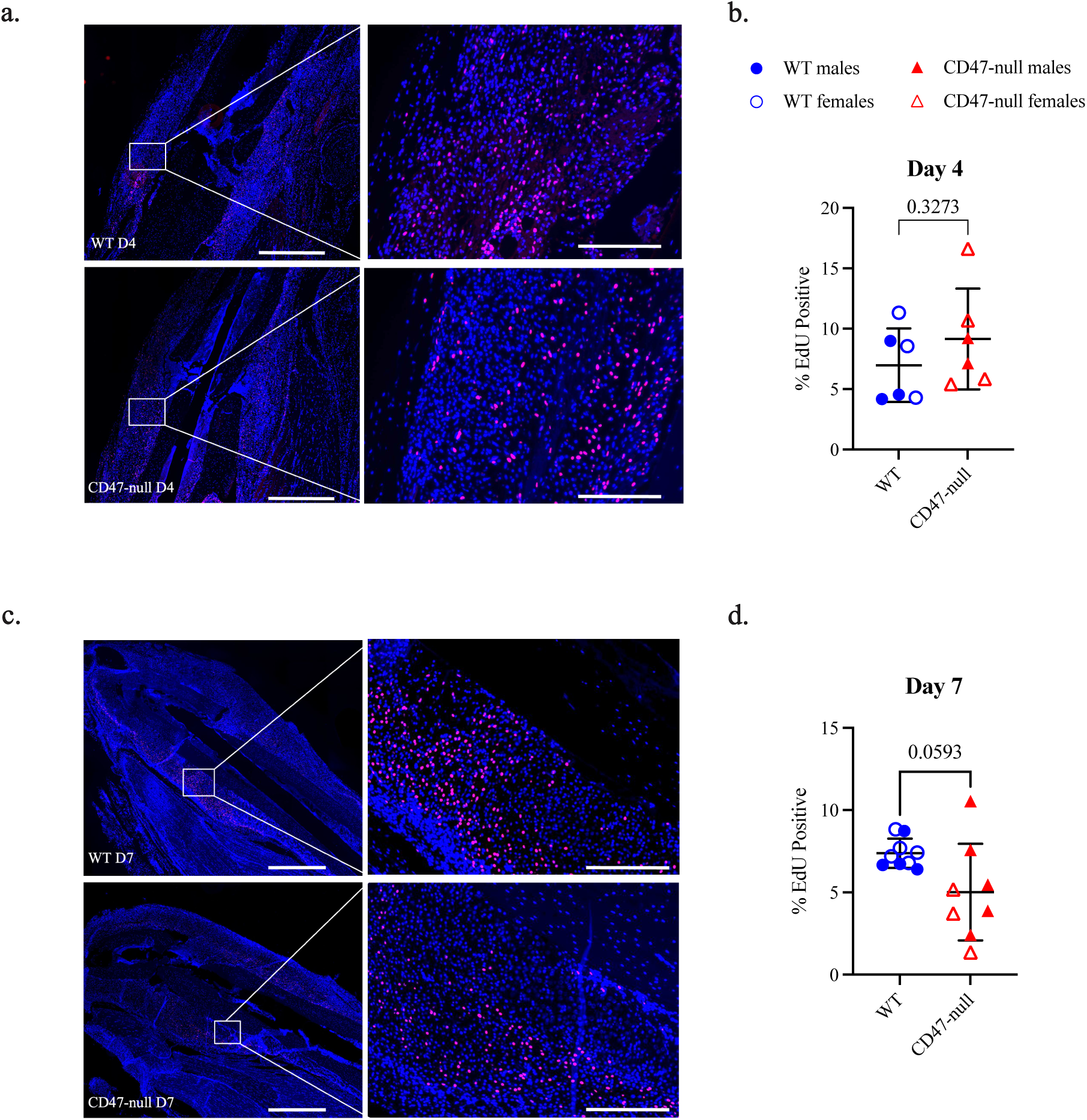
Genetic knockout of CD47 reduces cell proliferation in the early fracture callus. EdU expression at 4 and 7 days post ischemic tibia fracture in WT (n=6-9) and CD47-null (n=6-8) mice. **a,** Representative images of WT and CD47-null fracture calluses at day 4 post-fracture. **b,** % EdU positive cells in WT and CD47-null fracture callus at 4 days post-fracture. **c,** Representative images of WT and CD47-null fracture callus at day 7 post-fracture. **d,** % EdU positive cells in WT and CD47-null fracture calluses at 7 days post-fracture. Two-sided t-test. 10X stitched images, scale bar = 1000 µm. 10X zoomed images, scale bar = 200 µm.

### 2.13 Mice treated with *CD47* morpholino depict similar decreases in callus size

To investigate if the difference seen in the CD47-null mice ischemic fracture callus morphology was due to a developmental phenotype of complete CD47 knockout or due to an absence of CD47 signaling during the fracture healing process, a CD47 targeting morpholino ([M]CD47) that suppresses protein production was used to temporarily lower CD47 levels in mice. This was compared against a nonsense control morpholino ([M]Ctrl). Similar to the ischemic fracture callus observed in CD47-null mice, [M]CD47 treated WT mice demonstrated a reduction in early callus formation (**Figure 9**). At day 15 [M]CD47 mice developed a smaller and shorter callus with less bone and mineral content. However, the percent of bone in the smaller callus was higher, as was bone mineral density versus [M]Ctrl treated WT mice. This data demonstrates that the observed differences in CD47-null mice are not likely due to a developmental deficiency in CD47, but rather the effect of decreased CD47 in cells in callus tissue during fracture healing.

**Figure 9.**
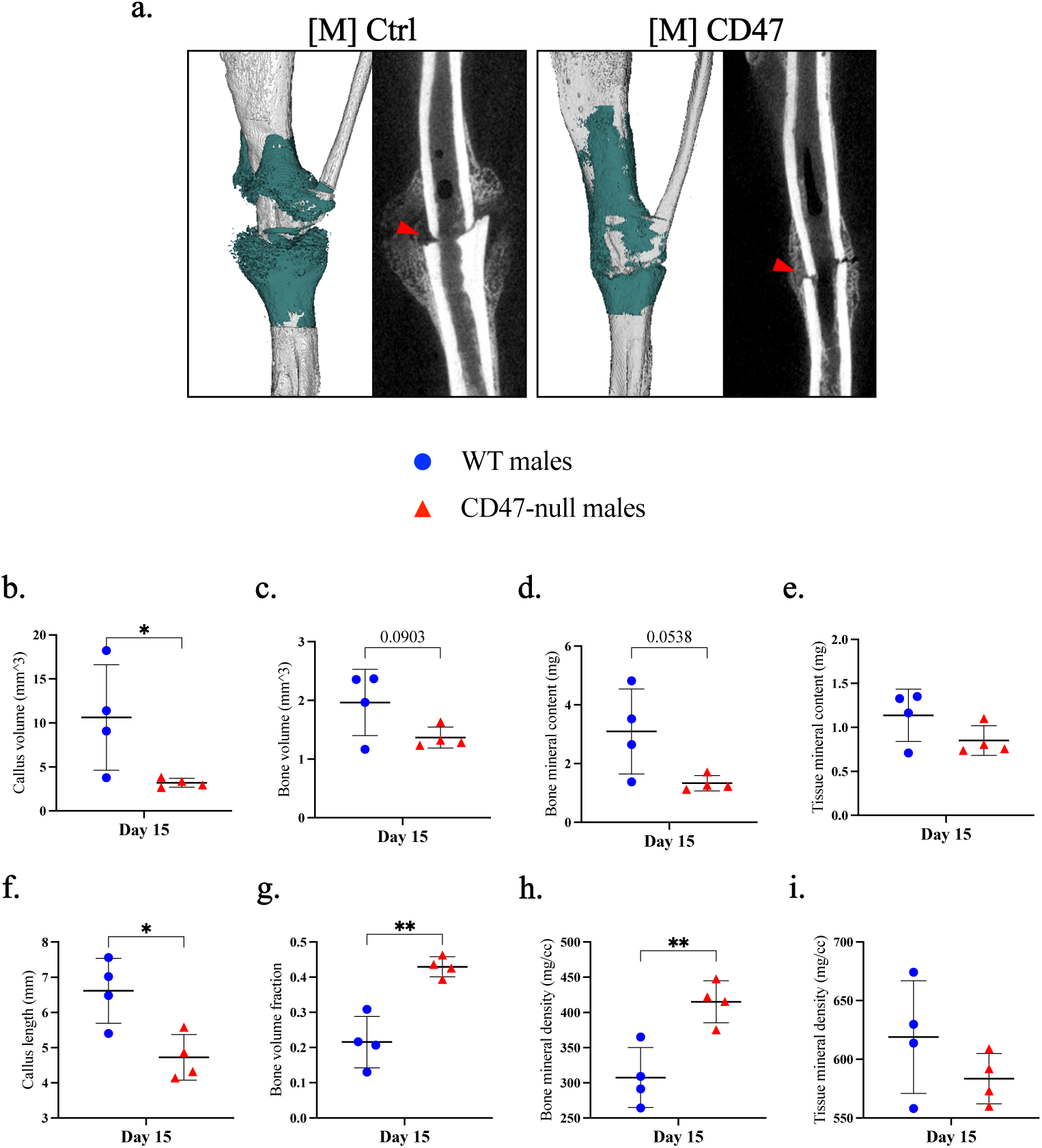
Disruption of CD47 using a morpholino inhibits early ischemic fracture callus formation. µCT analysis of ischemic tibia fracture callus of morpholino-control ([M]Ctrl) (n=4) and morpholino- CD47 ([M]CD47) (n=4) at day 15 post-fracture. **a**, 3D reconstructions (white background with teal mineralized callus highlight) and sagittal-plane reconstruction (black background) of [M]Ctrl (left) and [M]CD47 (right) mice at day 15 post-fracture. Location of fracture (red arrowhead) is marked on the sagittal reconstructions. **b-i**, Callus morphology (mean±SD) at day 15 post-fracture. \**P<*0.05, \*\**P<*0.01, two-sided t test.

### 2.14 Recovery of perfusion after induced ischemia in CD47 morpholino treated mice

To investigate the effect of temporary lowering of CD47 protein on recovery of ischemic injury, perfusion was measured prior to ischemia, immediately after, and at day 15 post-ischemia using laser doppler flowmetry of the plantar surface of the hindfoot. Relative change in perfusion was calculated as described. Similar to the decrease in perfusion immediately after ischemia seen in the WT and CD47-null mice, there was relative perfusion decrease of 88.42±1.39 and 87.61±4.46 in [M]CD47 and [M]Ctrl mice, respectively (**Supplemental Figure 7**). At day 15 post-ischemia, [M]CD47 and [M]Ctrl mice recovered to 37.894%±5.07 and 43.54%±6.14 of their baseline perfusion. Unlike the WT and CD47-null mice, there was no difference in perfusion recovery between [M]CD47 and [M]Ctrl mice. The lack of difference is likely due to the temporary nature of the change in CD47.

## 3 Discussion

The ubiquitously expressed protein CD47 has been shown to have both detrimental^13,27^ and protective effects on soft tissue repair^28^, however, the role of CD47 in mediating fracture repair has not been previously studied. Here we have demonstrated that CD47 deletion leads to diminished fracture callus size and bone formation, and this is accompanied by increased fibrotic tissue content. This is true when CD47 is deleted either constitutively or at the time of fracture using a morpholino.

A lack of CD47 negatively impacts callus cell proliferation at 7 days post-fracture. Notably by 7 days post-fracture, the inflammatory response has mostly resolved and there is considerable MSC expansion under typical healing conditions^29^ suggesting that the diminished proliferative effect we observed under CD47-null conditions is in the MSC. After expansion, MSC differentiate to become bone forming osteoblasts and cartilage forming chondrocytes. Reduced proliferation of both bone marrow-derived MSC as well as periosteal-derived MSC further corroborates dysfunctional expansion of the mesenchymal progenitor population. These results suggest that the phenotype of reduced callus formation may be driven by impaired mesenchymal cell proliferation. Multiple cancer studies have demonstrated the ability of CD47 to promote cell growth and migration through the MAPK/Erk signaling pathway. ^30–32^ Similar to the explosive outgrowth of cells in tumor growth, fracture callus formation requires a large expansion of mesenchymal cells. CD47 may similarly play a role in promoting rapid proliferation in the fracture callus through activating the MAPK/Erk signaling pathway as well. Decreased CD47-MAPK-Erk signaling could further explain the reduced osteoblastic differentiation and mineralization previously reported by other groups in the absence of CD47. ^33,34^

Interestingly, interrogation of the skeletal stem cell populations present in the uninjured periosteum showed no difference in baseline progenitor populations. This suggests a specific role of CD47 for progenitor expansion and differentiation post-injury. Our findings align with work by others that highlight the importance of CD47 not in uninjured tissues, but in facilitating wound repair. ^28^

Previous studies have suggested that inhibition of the CD47-TSP1 signaling axis results in improvement of soft tissue ischemic injuries. ^35^ To interrogate whether CD47 deletion could rescue ischemic fracture repair, we performed ischemic tibia fractures in CD47-null mice. In accordance with previous literature detailing an anti-angiogenic effect of CD47-TSP1 interactions, we did observe increased endothelial density early post-fracture (7 days post-fracture). However, we noted that despite the increased number of endothelial cells, we did not observe differences in distal limb perfusion, and there was reduced healing of the ischemic fracture callus under CD47-null conditions. This suggests that while inhibition of CD47 may improve callus vascularization, the deleterious effect of a global deletion on other aspects of healing, specifically stromal cell expansion, outweighs the positive effect this may present on healing in the context of ischemia. Targeted deletion of CD47 in endothelial cells may present a potential approach to improve vascularization.

While we have proposed potential mechanisms of action for CD47, we acknowledge CD47 has many different roles that could result in the diminished fracture callus phenotype observed. CD47 plays an important role in homeostasis of the immune system. Blockade of CD47 binding to macrophage SIRPα promotes phagocytosis. ^36,37^ CD47-SIRPα-signaling has also been shown to affect osteoclastogenesis. ^33^ Additionally, CD47 binds several integrins including α_v_β_3_, α_2_β_1,_ α_4_β_2_ to regulate cell functions such as migration, adhesion, and extracellular matrix organization. ^38^ Cell- and lineage-specific CD47-deletion experiments should be undertaken in the future to delineate if the CD47 callus phenotype is an indirect immunologically driven effect or a result of direct mesenchymal cell autonomous effects.

## 4 Materials and Methods

### 4.1 Study design and animal use

The study design was developed under the guidance of a university-wide AAALAC-accredited laboratory animal medicine program directed by veterinarians specialized in laboratory animal medicine. The protocol was reviewed and approved by the University of Michigan Institutional Animal Care and Use Committee (IACUC) and adhered to all applicable federal, state, local, and institutional laws and guidelines. CD47-null mice were developed by Lindberg et al. by replacing exon 2 of the Iap (*CD47*) genomic DNA with the neomycin resistance gene driven by the TK-promoter^39^ and were acquired from Dr. Jeff Isenberg. Knockout mice were back-crossed onto a C57Bl/6J background and compared against WT (WT) mice C57Bl/6J obtained from Jackson Labs (Bar Harbor, ME). All mice were bred in the same vivarium to ensure similar microbiomes and genotyped either in house using a polymerase chain reaction (PCR) with primers specific to CD47-null (forward: GGCATTGCCTCTTTGAAAATGGATA, reverse: TGGCTTCTGAGGCGGAAAG) knockout mice or by a third-party service (Transnetyx, Cordova, TN). Animals were socially housed and allowed ad libitum access to food and water. Euthanasia was performed in accordance with current American Veterinary Medical Association guidelines using CO_2_ and confirming death secondarily with the generation of a bilateral pneumothorax. For all surgeries, animals were sedated through inhalant anesthetics (4-5% isoflurane for induction; 1-2% for maintenance). Adequate sedation was confirmed by lack of responsiveness to a hind-limb toe pinch. The first dose of postoperative analgesia buprenorphine (0.1 mg/kg subcutaneously) was administered at the time of surgery. Ocular lubricant was applied to prevent corneal drying. Surgeries and quantitative analyses were conducted with investigators blinded from timepoint, genotype and treatment. For μCT and histology analysis, 6-14 animals were used per time point with a distribution of males and females. *In vitro* marrow-derived MSC assays were repeated 2-3 separate times with pooled cells from three mice. *In vitro* periosteum assays were performed in three separate experiments, each replicate is one mouse.

### 4.2 Femoral Fracture

Fractures were performed at 14 weeks of age during the late skeletal growth phase and just prior to peak bone mass. ^40,41^ The stabilization and limb fracture technique is published ^42^. Briefly, the femur was stabilized with a 23-gauge needle placed percutaneously into the medullary cavity. Radiographs were acquired to confirm that the pin extended the length of the medullary canal but did not protrude from the proximal or distal end of the femur. The limb was positioned prone in the fracture apparatus using a jig to center the point of impact at the middle third of the femur. A 272 gram weight was dropped from a height of 4.40 cm onto the femur and was stopped at an impact depth of 0.11 cm. Post-fracture radiographs were acquired to confirm the fracture location and pattern.

### 4.3 Hindlimb ischemia (HLI)

Hair was removed on one leg from mid-abdomen to mid-tibia, exposing the medial leg and knee joint, using depilatory cream. Mice were placed supine on the surgical field and the operative limb was secured in an abducted, externally rotated position. The surgical area was prepared and cleaned with two swabs of povidone-iodine and 70% EtOH and then protected with sterile drapes. A medial 15 mm incision was made proximal to the knee along the longitudinal axis of the limb. Retractors were placed and blunt dissection was used to visualize the femoral neurovascular bundle. The vein and nerve were dissected from the femoral artery to preserve vascular return and limb sensory-motor function. A 2 mm section of the femoral artery distal to the epigastric artery and proximal to the medial genicular artery was ligated using 8-0 nylon sutures and resected to induce deep distal acute ischemia. ^43,44^ The incision was closed using simple interrupted sutures with 6-0 nylon. Laser doppler measurements of hindlimb perfusion were used to confirm ischemia.

### 4.4 Laser Doppler

Hindlimb perfusion was measured using laser doppler blood flow monitor (Moor, moorVMS-LDF) equipped with a non-invasive skin probe (Moor, VP2). Animals were sedated as described above and maintained a respiratory rate of 55-100 breaths/min. Animals were placed supine on a heating pad (Hallowell EMC, 2789B hard pad & heat therapy pump) at 37.5 °C and remained undisturbed for 5 minutes to normalize to conditions. The doppler probe was placed on the plantar surface of the hindlimb with the distal boarder of the probe abutting the digital walking pads. Alternating perfusion measurements were taken three times for each limb and averaged.

### 4.5 Tibia Fracture with HLI

An ischemic tibial fracture was created by resecting the femoral artery, reducing distal perfusion as described above. The limb was stabilized using a 30-gauge intramedullary pin and a simple transverse fracture was created. Perfusion was tracked using laser doppler flowmetry and fracture pattern was confirmed with digital radiographs.

### 4.6 CD47 Morpholino Treatment

Expression of CD47 was disrupted using a translation-blocking antisense morpholino oligonucleotide. On days 2 and 5 post-fracture, mice were injected intraperitoneally using a 23-gauge needle with 1.0 nmol/g of morpholino in 750 μl of saline. Mice were injected with either a CD47 antisense vivo-morpholino (CGTCACAGGCAGGACCCACTGCCCA) or a vivo-morpholino standard control (CCTCTTACCTCAGTTACAATTTATA) oligonucleotide (Gene Tools, Philomath, OR).

### 4.7 Sample Preparation and Micro-computed Tomography (μCT)

Femurs were harvested at day 10 and 20 post-fracture for non-ischemic injury experiments. Tibia were harvested at day 10, 15, and 20 for knockout ischemic studies and at day 15 post-fracture for morpholino ischemic experiments. Bones were fixed in 4% paraformaldehyde (PFA) for 24 hours at 4 °C under gentle agitation. After fixation, the intramedullary pins were carefully removed without disrupting the callus by extracting them from the distal end of the femur using small-nosed needle nose plyers. Femurs were placed in a 4-limb positioning jig and scanned in eXplore Locus scanner (GE Healthcare). The femur containing jig was immersed in water and specimens were scanned at an 18 µm voxel size using the following settings: 80 kV, 80 µA, 1600 ms, 400 views. Reconstruction and analysis were performed in MicroView (Parallax Innovations). A callus region of interest (ROI) was created by manually sectioning around the perimeter of the callus every 10 slices and then performing a spline interpolation between each individual section. Cortical bone was subtracted from the callus ROI using a similar sectioning and interpolation technique. For day 10 fractures, a 3.00 x 3.00 x 5.00 cylindrical ROI was centered around the fracture to avoid sectioning errors caused by vague delineation of density between callus edge and soft tissue. The callus ROI was analyzed for callus length, callus volume (commonly referred to as total volume (TV)), bone volume (BV), total mineral content (TMC), and bone mineral content (BMC) using a threshold of 1650 Hounsfield units (HU). The threshold was established by averaging the bone threshold of 10 whole intact femurs using the automatic bone density algorithm MicroView based on thresholding methods. ^45^ Bone volume fraction (BVF) was calculated by dividing the BV by the TV and represents how much of the callus contains mineralized tissue. Tissue mineral density (TMD) was calculated by dividing the TMC by the BV, and bone mineral density (BMD) was calculated by dividing BMC by TV. Representative images were created using isosurfaces of cortical bone and callus with a threshold set at 1200 HU. Isosurfaces were post-processed using smoothing.

### 4.8 Sample Preparation for Histology and Immunofluorescence

Tibiae were harvested at days 4 and 7 post-fracture for knockout experiments and fixed in 4% paraformaldehyde for 24 hours under gentle agitation. After fixation, the intramedullary pins were carefully removed without disrupting the callus by extracting them from the distal end of the tibia using small-nosed needle plyers. Tissues were formalin-fixed, paraffin-embedded using a tissue processor (Leica ASP 300S) and sectioned in the sagittal plane at 5 for immunofluorescent staining or 10 µm sections for Safranin O analyses. Slides were deparaffinized using a sequence of xylene and ethanol dilutions.

#### 4.8.1 Histomorphometry Analysis

Sagittal sections (10 µm) through the entire block were cut and every thirtieth slide was stained with Safranin-O/fast green. Adjacent slides were stained with modified Milligan’s Trichrome. An Olympus CAST microscope (Center Valley, PA) and software made by Visiopharm (Hørsholm, Denmark) was used to perform stereology. ^46^ Histomorphometry was used to calculate callus volume and the volume of cartilage, bone, fibrous tissue, and marrow within the comprising callus. Total volumes were estimated using Cavalieri’s principle for a conical frustum as in the past. ^47^

#### 4.8.2 EdU Analysis and Quantification

Mice were injected with 10mM EdU intraperitoneally at 4 hours prior to harvest at day 4 and day 7. Tissue samples were harvested and prepared as described above. Sagittal sections (10 µm) were utilized for EdU quantification. Post-deparaffinization, slides were permeabilized with 0.5% Triton X-100 for 10 minutes. A commercially available EdU Click-it Kit (Fisher, C10339) was used to label EdU positive cells. Slides were washed 3 times in PBS and counter stained with DAPI. Entire sections were imaged at 10X resolution and stitched in both Texas Red and DAPI channels. Photoshop was used to define the callus ROI and CellProfiler was used to quantify EdU positive nuclei within the ROI using Texas Red. DAPI was used to quantify nuclei. These were overlayed to determine the percent of EdU positive nuclei for each callus.

#### 4.8.3 Endothelial Cell Density Quantification

Heat-mediated antigen retrieval was performed, and tissues were permeabilized with 0.5% Triton-X 100 in phosphate buffered saline (PBS) prior to immunofluorescent staining. Immunofluorescence staining was performed on 5 µm sagittal sections using rabbit anti-mouse CD31(Abcam, ab281583) and rat anti-mouse Emcn primary antibodies to label endothelial cells, followed by AF750 and AF594 secondary antibodies. DAPI stain was used to identify nuclei. Primary, secondary, isotype, and unstained controls were used to detect non-specific staining or autofluorescence. 2-4 images at the central callus and distal callus regions were taken using an automatic microscope (Agilent, BioTek Lionheart FX, Santa Clara, CA). Central callus regions were defined as proximal to the fracture site on either side of the bone, and distal callus was defined as the periphery of the fracture callus. The Vessel Analysis plug-in on ImageJ was used to calculate endothelial cell density in central and distal regions of the fracture callus.

### 4.9 Marrow Cell Harvest

Femurs and tibias were harvested from adult mice and dissected of all soft tissue under sterile conditions. The distal femur and proximal tibia were separated at the level of the metaphysis to expose the medullary canal. The long bones were placed exposed end down in a 0.5 ml microcentrifuge tube with a hole in the bottom made with an 18-gauge needle. The microcentrifuge tube was nested in a 1.5 ml microcentrifuge tube and centrifuged at 10,000 x g for 15 minutes. Cells were suspended in culture media and passed through a 70 μm strainer. Cells were then pelleted and washed twice with culture media.

### 4.10 Periosteum Cell Harvest

Femurs and tibias were harvested from adult CD47-null and C57 Bl/6 mice. The bone was isolated from the muscle and the articular and epiphyseal ends were dipped into pre-warmed 5% agarose gel. After solidification of the agarose, the femurs and tibias were placed in a 10% Collagenase P Digestion buffer at 37 °C with agitation. After 10 minutes, the buffer was discarded to remove contaminating cells. Bones were transferred to a new tube containing 10% Collagenase P Digestion for 1 hour at 37 °C with agitation. Cells were passed through a 70 μm strainer, pelleted and resuspended in culture media.

### 4.11 Endothelial Cell Harvest and Purification

Endothelial cells were isolated from the lungs of 4-week-old mice. Lungs were removed using sterile scissors and cut into <0.2 cm pieces. Lung tissue was incubated under agitation at 37 °C in a 50 ml conical tube with 10 ml of collagenase A in PBS (5 μg/ml). After 30 minutes, 20 ml of PBS was added to the conical tube and then vigorously shaken and to further dissociate tissue. The cell suspension was filtered through a 70 μm strainer, pelleted, and washed once with endothelial cell media (Lifeline Cell Technology, VascuLife VEGF Endothelial Complete, Frederick, MD). The cell suspension was sorted for CD31+ endothelial cells using anti-CD31 MicroBeads (Miltenyi Biotec, 130-097-418; San Diego, CA) and a magnetic cell isolation system (Miltenyi Biotec, QuadroMACS Separator & LS Columns). CD31+ cells were cultured in endothelial cell media on 2% gelatin coated plates and flasks or used immediately for flow cytometry.

### 4.12 Colony Forming Unit – Fibroblast (CFU-F)

Cells harvested from marrow were plated at a density of 2x10^5^ cells/cm^2^ in 60 mm dishes and grown in 5% CO_2_ at 37 °C. Half of the culture media (Alpha-MEM, 10% FBS, 1%, L-glutamine, 1% anti-anti) was exchanged every three days until cell harvest. At day 12 post-plating, bone marrow-derived plates were stained with alkaline phosphatase to mark mesenchymal stem cells. Dishes were washed twice with PBS and then fixed with Citrate-Acetone-36% Formaldehyde (ratio, 25:65:8) for 30 seconds and then rinsed with deionized water. Naphthol AS-BI Alkaline Solution (Sigma-Aldrich, 861-10) was added to the plates at 25 °C for 15 minutes and then rinsed with deionized water. Plates were counterstained at 25 °C for 2 minutes with Neutral Red (Sigma-Aldrich, N6264). Finally, dishes were rinsed with tap water and allowed to air dry. Individual dishes were examined using an upright brightfield stereo microscope (Bausch & Lomb) placed over a 1 cm grid. Groups of 25-100 ALP positive cells were counted as small colonies and confluent groups of >100 ALP positive cells were counted as large colonies. Counting was performed blinded to cell genotype. Cells harvested from the periosteum were plated at a density of 1x10^5^ cells/cm^2^ in 60 mm dishes and grown in 5% CO_2_ at 37 °C. At day 10, post-plating alkaline phosphatase staining was performed on periosteal MSC as described above. Individual dishes were imaged in color brightfield (BioTek, Lionheart FX) to analyze and count clusters with alkaline phosphatase staining.

### 4.13 Cellular Proliferation (MTT)

Metabolically active cells were quantified using an MTT (3-(4,5-Dimethylthiazol-2-yl)-2,5-diphenyltetrazolium Bromide) assay which measures formazan produced by cleavage of tetrazolium salts (MTT) by the succinate-tetrazolium reductase system. A commercial cell proliferation kit (Millipore Sigma, Cell Proliferation Kit I (MTT)) was used for mesenchymal stem cells and endothelial cells. Cells were grown in 12-well plates in 5% CO_2_ at 37 °C as described. On the day of assay, MTT was added to the cells (10 μl MTT per 100 μl of media). Cells were returned to the incubator for 4 hours and then a solubilization solution was added to the cells (100 μl solubilization solution per 100 μl of media). Cells were incubated overnight to solubilize the purple formazan crystals. The next day 210 μl from each well was added in triplicate to a 96-well plate. Formazan concentration was quantified on a multi-mode microplate reader (Molecular Devices, SpectraMax M3) using a 575 nm wavelength for formazan spectrophotometric absorbance and 650 nm for reference. Readings were averaged across technical replicates.

### 4.14 Cellular Proliferation (CCK8)

Metabolic activity within periosteal MSC was quantified using a Cell Counting Kit 8 assay (WST-8 / CCK8; Abcam, Cambridge, UK). Water soluble tetrazolium salts are used to quantify cell viability through the production of formazan dye similar to that of an MTT assay. Periosteal MSC were isolated and grown in 12 well plates at 5x10^4^ cells/well. Starting at day 1, every other day CCK8 reagent was added to the cells every other day for 9 days. The plates were returned to the incubator and analyzed 2 hours later. 100 μl from each well was added to a 96 well plate and read on the spectrophotometric at 460 nm. Readings were averaged across technical replicates.

### 4.15 RNA isolation and gene expression (quantitative RT-qPCR)

mRNA was isolated from adherent cells using 4 ml TRIzol and extracted using acid guanidinium thiocyanate-phenol-chloroform with GlycoBlue co-precipitant. mRNA was purified with Qiagen RNeasy Midi spin columns (Qiagen, Redwood City, CA) with on-column DNase digestion per the manufacturer’s protocol. cDNA was reverse transcribed from mRNA with Superscript III (Invitrogen, ThermoFisher) and quantitative RT-PCR reactions were conducted with 20 ng of template using custom primers (**Table 1**) and SYBR Select master mix (Applied Biosystems, Waltham, MA) for 50 cycles. Gene expression was calculated by normalizing to GAPDH (ΔCT) and then then to WT mice (ΔΔCT). Fold change was calculated as 2^−ΔΔCT^.

**Table 1:**
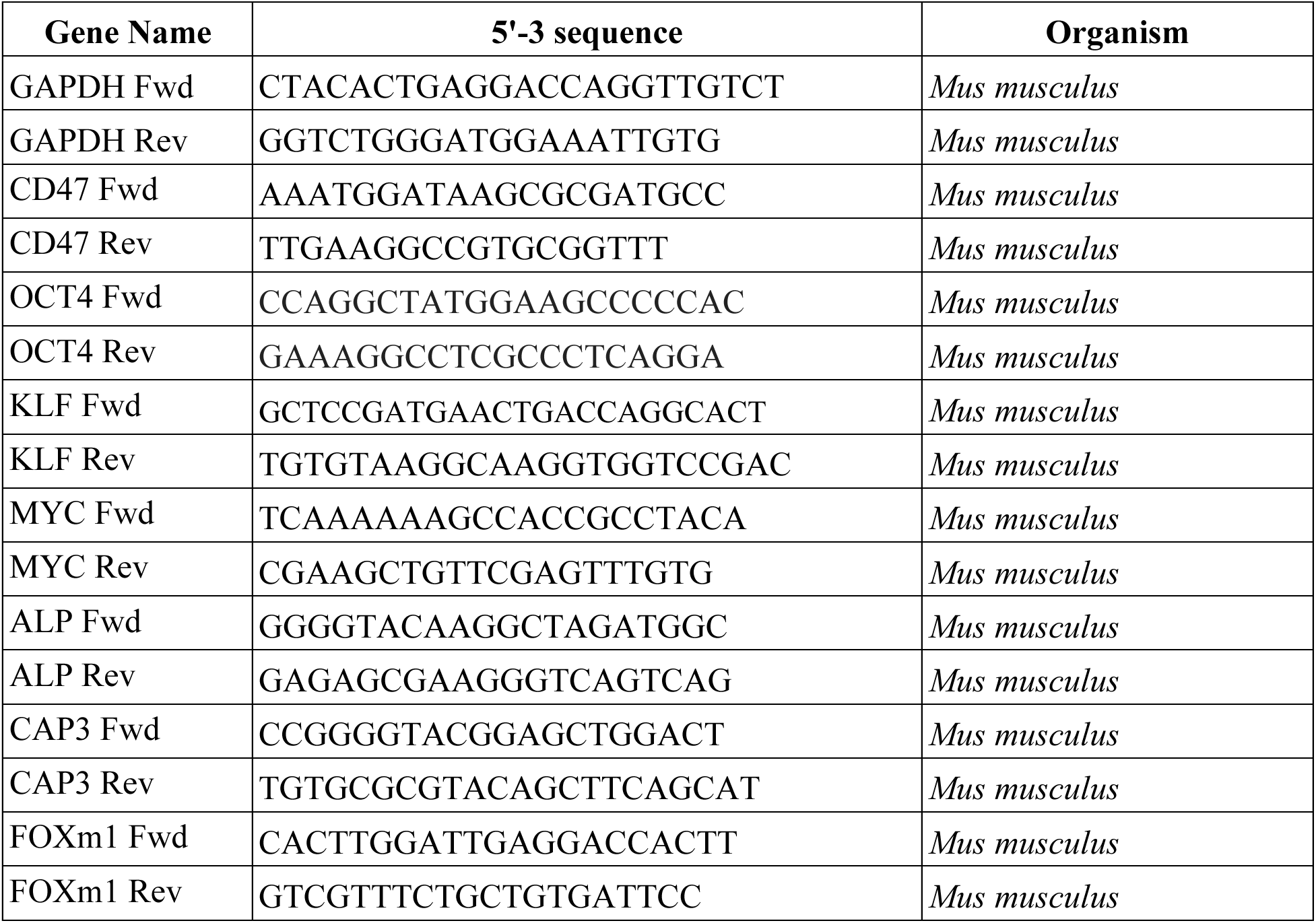
Sequences of reverse and forward primers used to assess RNA through q-PCR analysis.

### 4.16 Flow Cytometry

#### 4.16.1 Cell Cycle and Apoptosis Analysis

Cells for flow cytometry were counted and 5x10^5^-1x10^6^ cells were placed in individual 12 x 75 mm round-bottom tubes suspended in flow cytometry staining buffer (Invitrogen, 00-4222-26). For apoptosis, a Caspase-3/7 assay kit was used (Invitrogen, C10427). Each tube was brought to a volume of 1 ml staining buffer and 1 μl of CellEvent Caspase-3/7 reagent was added. Cells were incubated at 37 °C in the dark for 25 minutes and then 1 μl of SYTOX AADvanced dead stain solution was added. Cells were incubated at 37 °C for 5 minutes and then analyzed. For cell cycle analysis, cells were fixed with 0.5 ml of 100% cold ethanol for 20 minutes at 4 °C under gentle rotation. Cells were pelleted and the ethanol was decanted. Cells were resuspended in 1 ml of staining buffer and 4 drops (164 μl) of Propidium Iodide Ready Flow (Invitrogen, R37169) was added to the cells and incubated at 25 °C for 20 minutes. Cells were analyzed on a ZE5 Cell Analyzer (Bio-Rad) and data processed using FCS Express 6 Flow Cytometry software (De Novo Software, Pasadena, CA). The percentage of the cell population in G1, or S/G2 phase of the cell cycle was calculated by fitting Propidium Iodide excitation counts using FCS Express: Multicycle (De Novo Software).

#### 4.16.2 Endothelial Cell Immunophenotype

Endothelial cells were harvested from lung tissue and sorted as described. Cells for flow cytometry were counted and 5x10^5^-1x10^6^ cells were placed in individual 12 x 75 mm round-bottom tubes suspended in flow cytometry staining buffer (Invitrogen, 00-4222-26). A conjugated antibody for CD31 and control (Invitrogen, 12-0311-82, preparation: conjugated PE, host: rat, isotype: IgG2a, clone: 390) were added to the cells at 4 °C for 1 hour. Cells were pelleted and washed twice with staining buffer and suspend 1 ml of staining buffer with DAPI for an end concentration of 1 x 10^6^ cells/ml and DAPI (0.2 ug/ml; BD Biosciences, 564907) added. Cells were analyzed on a ZE5 Cell Analyzer (Bio-Rad) and data was analyzed using FCS Express 6 Flow Cytometry (De Novo Software).

#### 4.16.3 Periosteal Skeletal Stem Cell Phenotype

Periosteal MSC were harvested as described from C57 Bl/6 and CD47-null mice. Post digestion, cells were resuspended in PBS and stained with a panel to mark skeletal stem cells^24,48^ using conjugated antibodies for CD90, Sca1, CD140a, CD51, CD200, Ly51, CD105, and a lineage negative cocktail. Samples were stained with fixable viability dye for 30 minutes. TrueStain FcX block (Biolegend, San Diego, CA; 156604) was added and cells were pelleted and washed prior to antibody staining at 4 °C for 1 hour. Following staining, samples were pelleted, washed, and fixed using commercially available Fixation/Permeabilization Buffer (Biolegend, 426803). Cells were analyzed on an LSRFortessa Cell Analyzer (BD), and data was analyzed using FlowJo (BD).

### 4.17 Osteogenic Differentiation

Marrow cells were cultured in a T75 flask for 7 days and then trypsinized, counted, passaged, and plated in osteogenic media (Alpha-MEM, 0.5% β-glycerophosphate, 0.1% ascorbic acid-2-phosphate, 1% L-glutamine, 1% antibiotic-antimycotic) at density of 200,000 cells per well in a 6-well plate. At day 14, cells were stained with ALP as described or Alizarin Red S (ARS) to image mineralization. For ARS staining, cells were washed gently with PBS and fixed with 4% PFA for 60 minutes under slow rotation. Filtered 1% Alizarin Red S (Millipore, A5533) was added to the cell monolayer for 15 minutes in the dark at 25°C. Plates were washed 3-4 times with distilled water until background staining was removed. ALP and ARS plates were imaged using an inverted conventional brightfield microscope (BioTek, Lionheart FX) affixed with RGB imaging cubes. Stitched imaged were acquired using a Phase Plan Fluorite 1.25x air objective, registered using the blue channel, and stitched with 10% overlap. Stitched images were analyzed for staining quantity using Fiji ImageJ2. ^49^ Only the blue channel was used for analysis. A 560 x 560-pixel circular ROI was created to capture the whole plate but exclude edge artifacts. Image thresholds were set to 21/124 and 139/158 for ALP and ARS, respectively. Images were then measured for percent of ROI with positive staining.

### 4.18 Statistical Analysis

The μCT, stereology, flow cytometry, and gene expression datasets were grouped by genotype as an independent variable (WT or CD47-null; [M]Ctrl or [M]CD47) and by day as indicated. Data were tested for normal distribution and equal variances before analysis. Mean and standard deviation (SD) were calculated for continuous variables. While sex of the sample is shown where applicable for fracture phenotype analyses using μCT, histomorphometry, and immunofluorescent analysis, sex was not considered in the statistics of this study. Categorical variables were expressed as number and percentage. Two-sided t-tests were performed to test for differences between genotype at each timepoint. A one-way ANOVA was performed to assess differences in perfusion between limbs and across genotype. Perfusion recovery data was fit to a one-phase non-linear curve (Prism 8, GraphPad, San Diego, CA) and compared using extra sum-of-squares F test. Data was aggregated and analyzed using open-source R^50^ and Prism 10. Statistical significance was set at P ≤0.05. Points of significance are annotated graphically in figures with \**P* <0.05, \*\**P* <0.01, and \*\*\**P* <0.001.

### Data Availability

Data available on request from the authors.

## Acknowledgements

Research reported in this manuscript was supported by the National Institute of Arthritis and Musculoskeletal and Skin Diseases (NIAMS) of the National Institutes of Health (NIH) under award numbers F30AR071201 (RLZ) and R01 AR066028 (KDH). Additional research support is provided by the NIH under a training award T32TR004371 (CAC). The content is solely the responsibility of the authors and does not necessarily represent the official views of the NIH.

