## Supplemental Figures for "CD47 is Required for Mesenchymal Progenitor Proliferation and Fracture Repair"

### Supplementary Figures

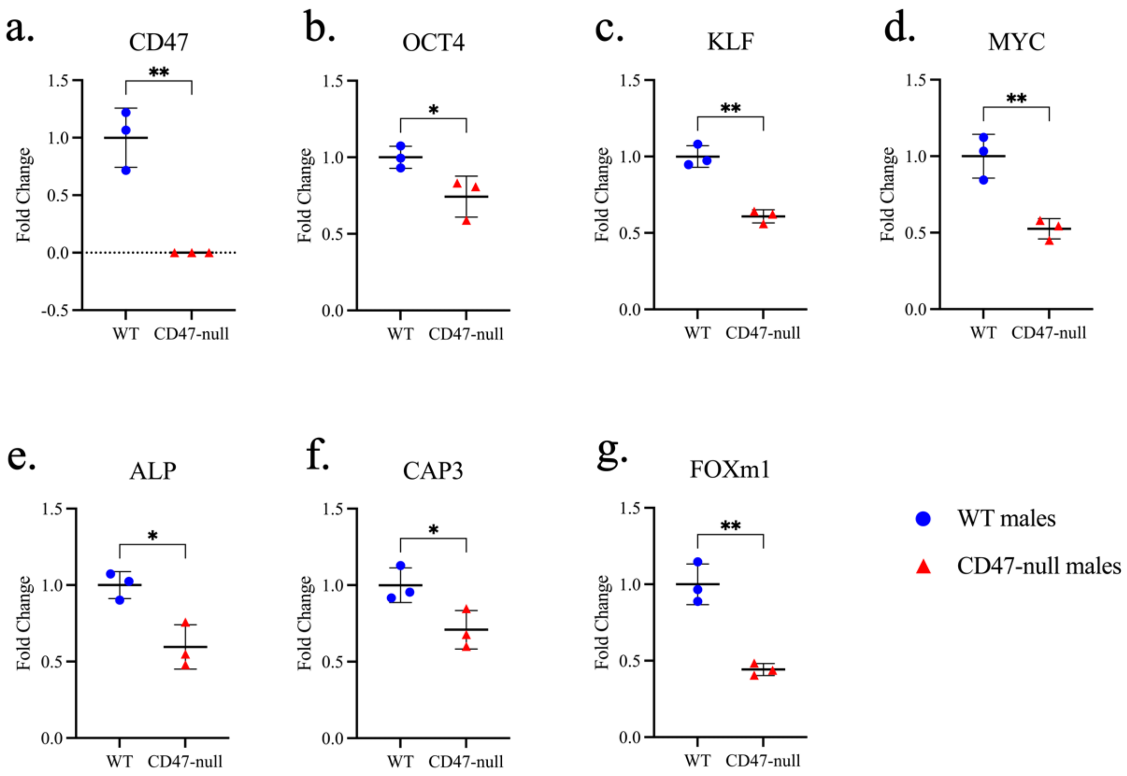


**Supplementary Figure 1: Loss of CD47 decreases expression of markers for pluripotency, proliferation, and apoptosis in MSC.** Gene analysis of marrow cells harvested from the femur and tibia of WT (n=3) and CD47-null (n=3) after 1 passage. **a-g**, Pluripotent stem cell markers, Oct4, Klf4, c-Myc, alkaline phosphatase (ALP), apoptosis marker caspase 3 (CAP3), and proliferation marker forkhead box M1 (FOXm1). Graphs indicate WT and CD47-null gene expression normalized to the housekeeping gene, GAPDH, relative to WT mice. Mean±SD, *$P$<0.05, **$P$<0.01, ***$P$<0.001, two-sided t tests.

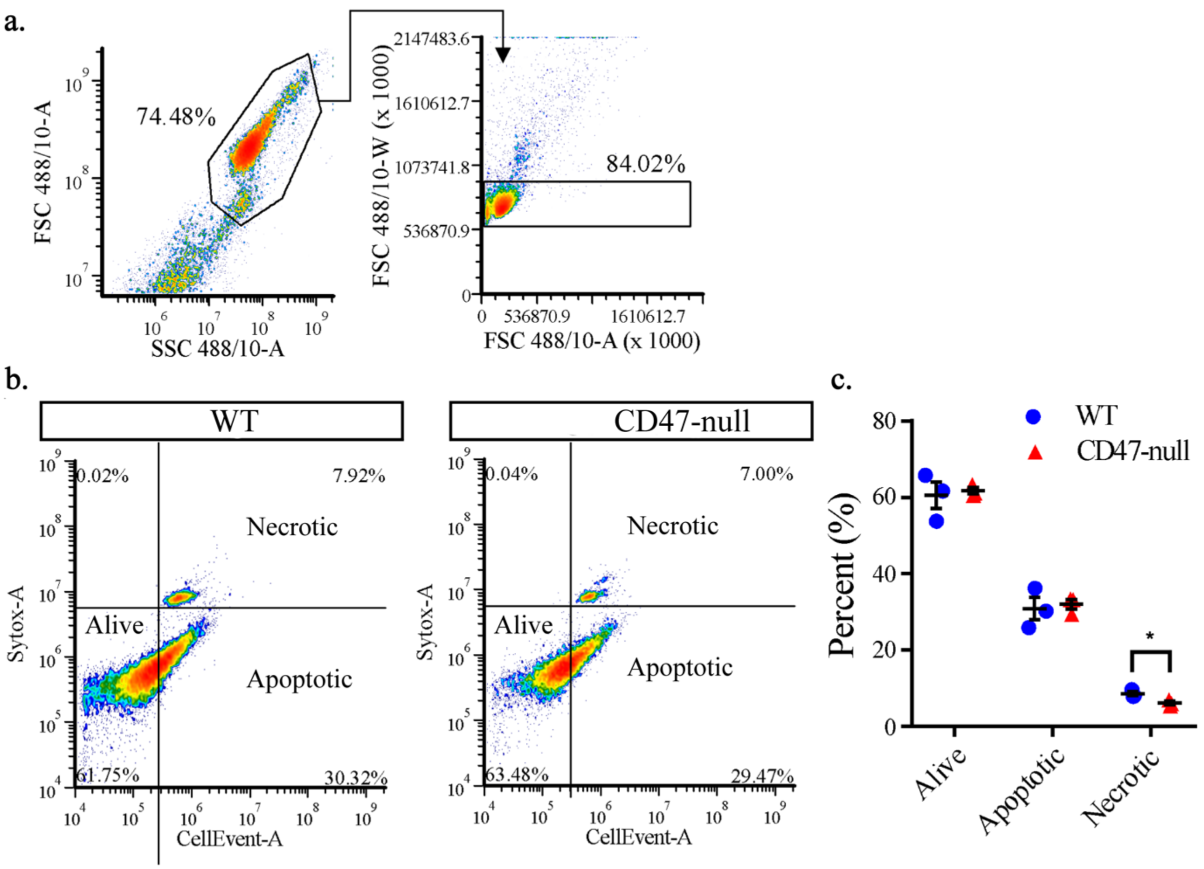


**Supplementary Figure 2: Loss of CD47 does not increase mesenchymal stem cell apoptosis.** Apoptotic analysis of cells harvested from the femur and tibia of WT (n=3) and CD47-null (n=3) mice. **a**, Flow cytometry gating scheme for cell density (left) and doublet discrimination (right). **b**, Representative plots of WT and CD47-null with quadrant overlay to segment alive, apoptotic, and apoptotic/necrotic cells. **c**, Comparison of WT and CD47-null population percentage of alive, apoptotic, and apoptotic/necrotic cells. Mean±SD, **P*<0.05, two-sided t test.

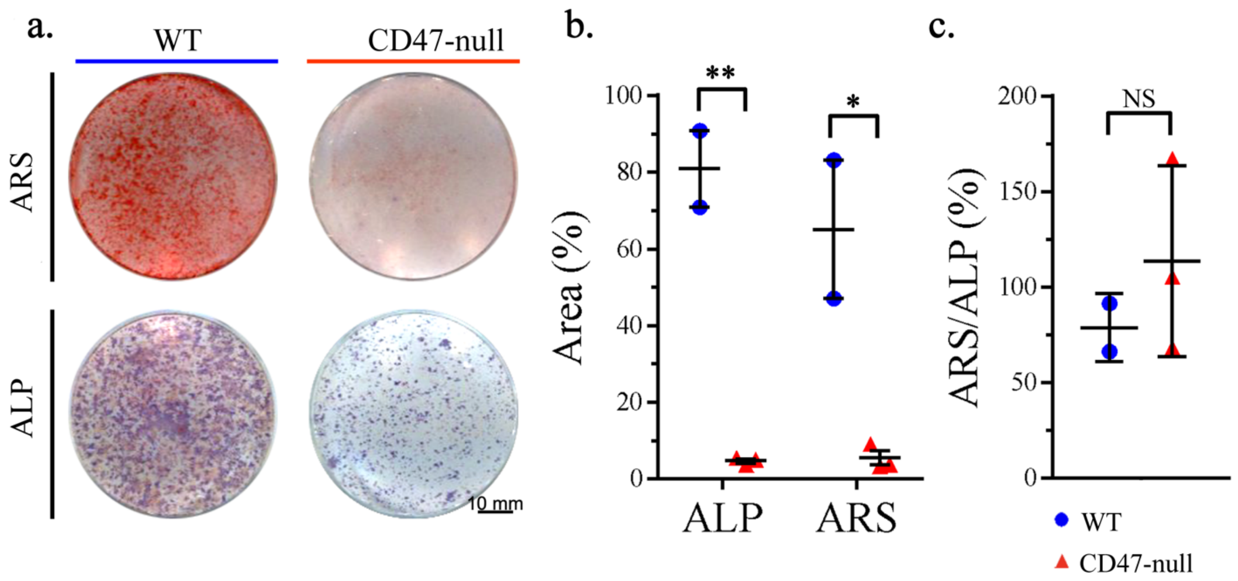


**Supplementary Figure 3: Loss of CD47 does not inhibit osteoblast differentiation.** Osteogenic analysis of marrow cells harvested from the femur and tibia of WT (n=2) and CD47-null (n=3) mice. **a**, At day 14, plates with stained with either alizarin red s (ARS) or ALP/Neutral Red. **b,** % Area of ALP and ARS staining. **c,** % area of ARS staining relative to ALP staining. Mean±SD, *$P<0.05$, **$P<0.01$, two-sided t test.

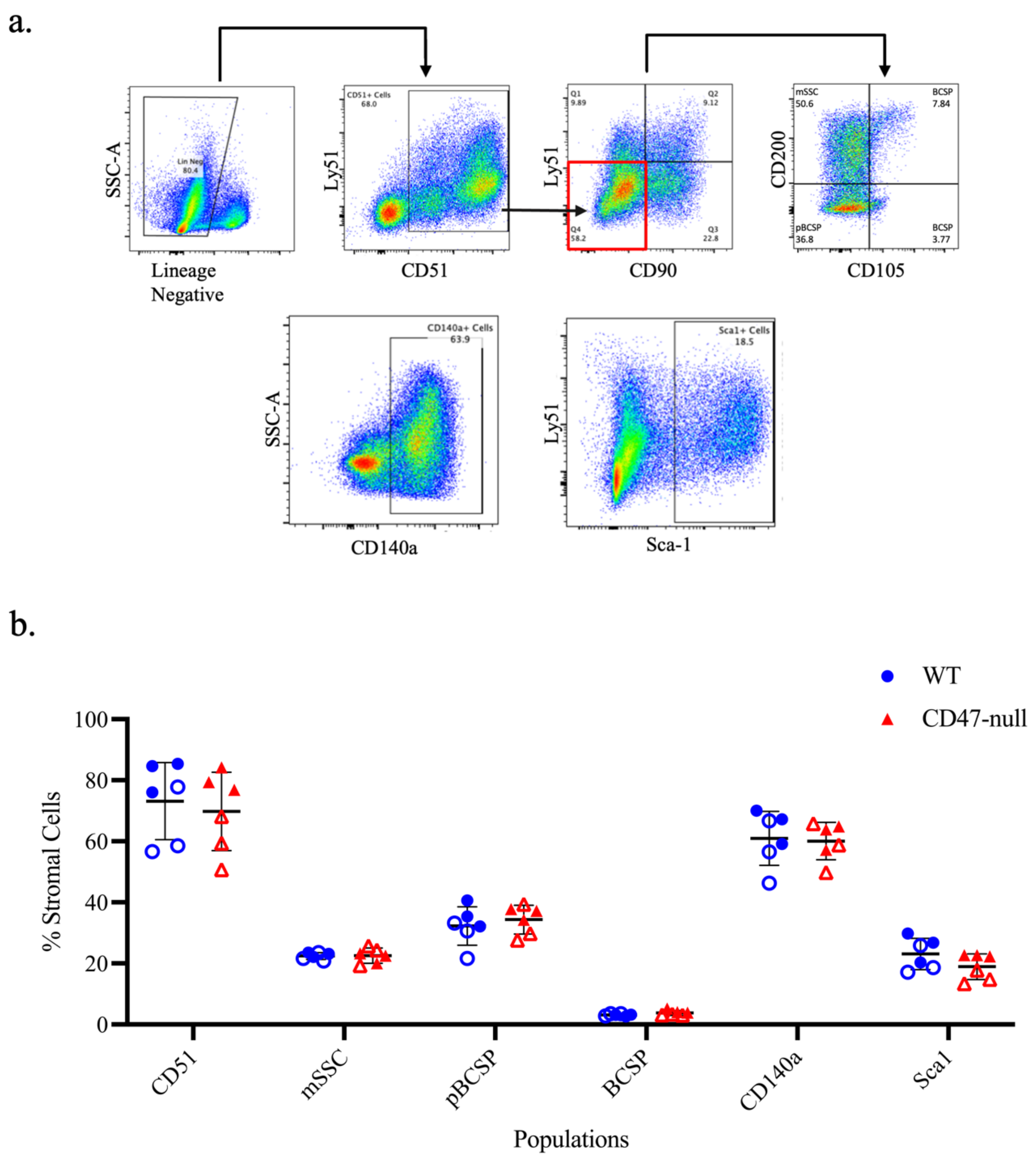


**Supplementary Figure 4: Loss of CD47 does not impact baseline skeletal stem cell numbers.** Skeletal stem cell marker analysis of periosteal MSC harvested from the femur and tibia of 10-week-old WT (n=6) and CD47-null (n=6) mice. Primary cells were stained with mesenchymal skeletal stem cell panel (fixable viability dye, lineage Negative, CD50, CD90, Ly51, CD105, CD200, CD140a, Sca1), fixed and analyzed for flow cytometry. **a**, Gating scheme for mesenchymal stromal stem cells (mSSC), pre-bone cartilage progenitor cells (pBCSP), bone cartilage progenitor cells (BCSP), CD140a+ cells, and Sca1+ cells. **b**, bar plot depicting % of total stromal cells (Lineage negative cells) that each population contributes. Mean±SD, two-sided t test.

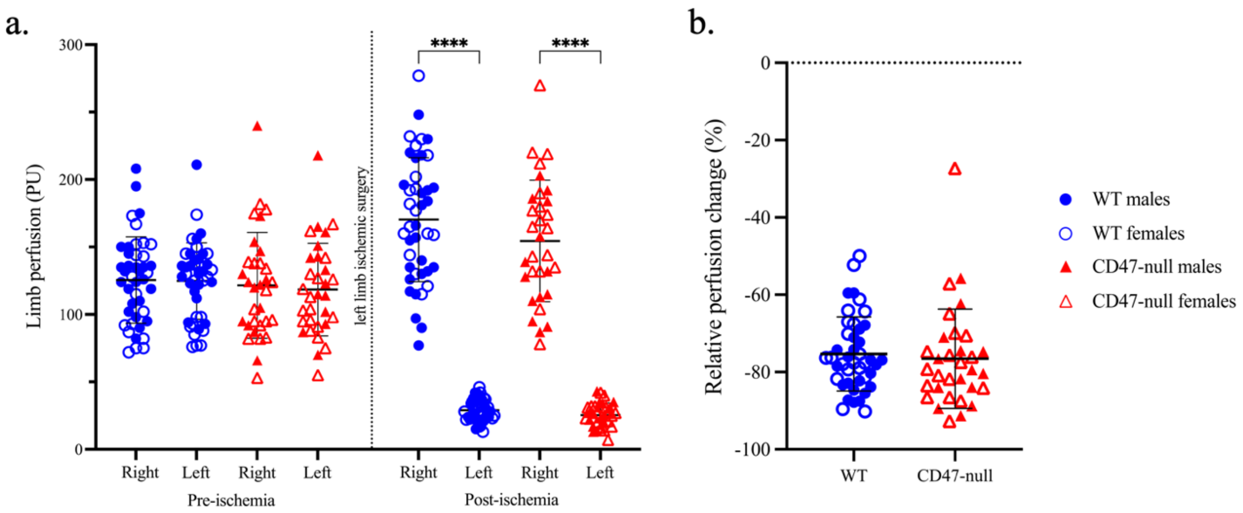


**Supplementary Figure 5: Genetic knockout of CD47 has no effect on baseline vascular function or immediate ischemic response.** Relative perfusion of WT (n=43) and CD47-null (n=32) mice before ischemia and 9 days post-ischemic tibia fracture using laser doppler flowmetry. **a**, Left: limb perfusion (mean±SD) pre-ischemic surgery for WT and CD47-null mice in right and left limb. No difference was seen between limb or genotype (one-way ANOVA). Right: limb perfusion (mean±SD) post-ischemic surgery for WT and CD47-null mice in right and left limb (One-way ANOVA). **b,** Relative perfusion change post-ischemia was determined by calculating the percent change in the right limb ratio from pre-ischemia to post-ischemic. WT and CD47-null mice ischemic limb perfusion decreased by -81.83%±1.03 and -82.04%±1.449, respectively. No difference was observed in relative perfusion, two-sided t test. ****$P$<0.0001, one-way ANOVA.

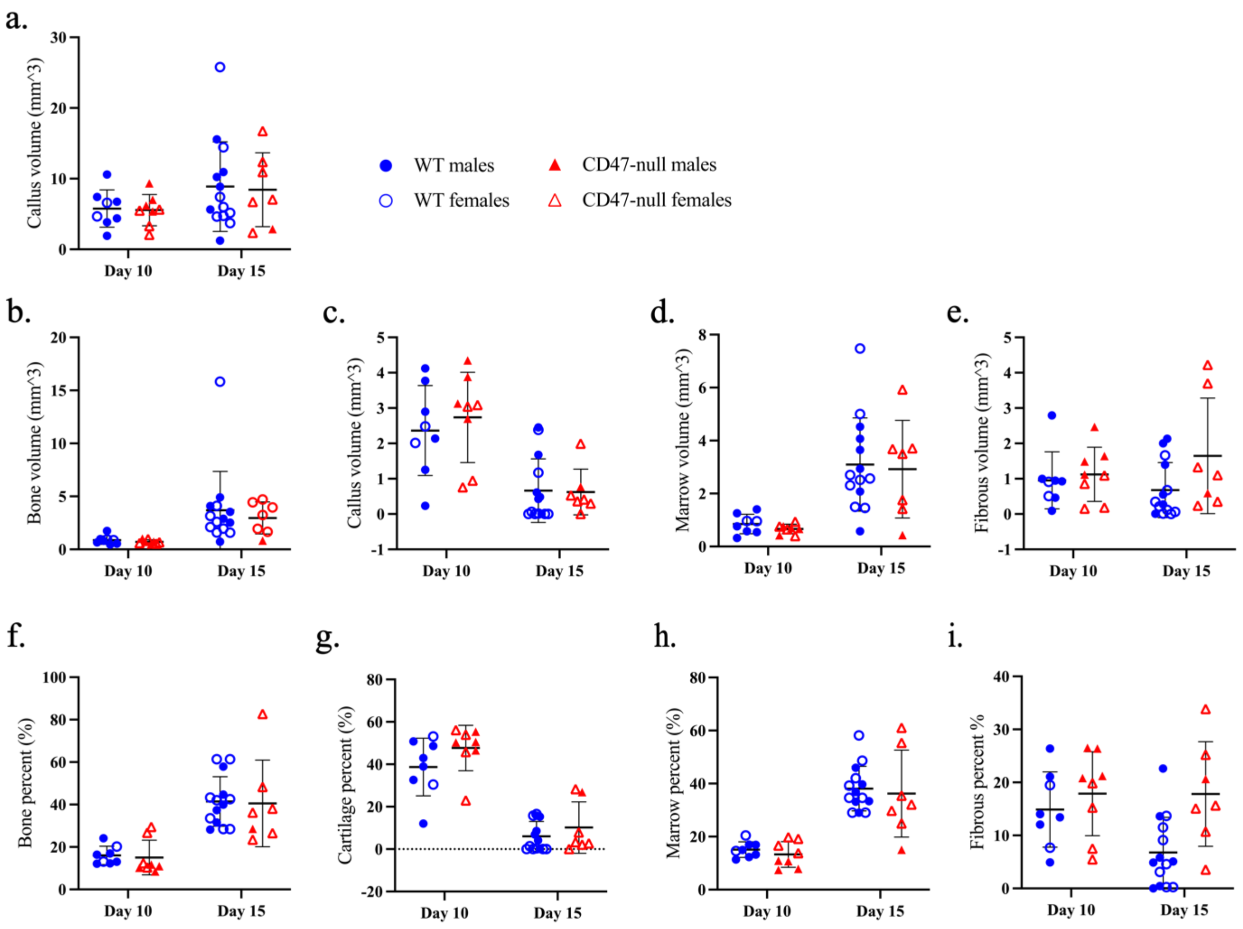


**Supplementary Figure 6: Histological data of CD47-null mice with ischemic fracture does not correlate with μCT data.** Histomorphometry of ischemic tibia fracture callus of WT (n=8-9) and CD47-null (n=6-8) mice at 10- and 15-days post-fracture. **a-i**, Callus composition (mean±SD) at day 10 and 15 post-fracture. Two-sided t test.

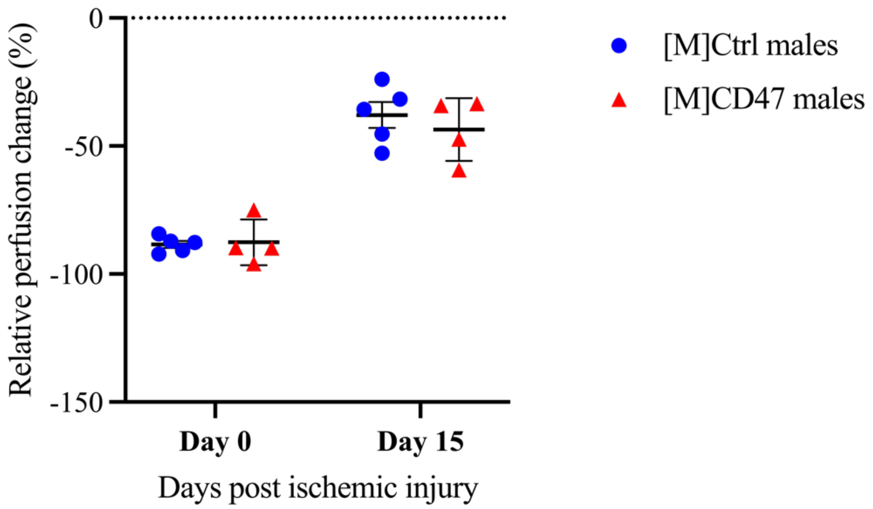


**Supplementary Figure 7: Disruption of CD47 using a morpholino does not limit recovery of perfusion after induced ischemia.** An ischemic tibial fracture was induced in WT mice prior to intraperitoneal injection with either morpholino-control ([M]Ctrl) (n=5) or morpholino-CD47 ([M]CD47) (n=4) at days 2 and 5 post fracture (1 nmol/kg). Perfusion measurements were performed prior to ischemia, immediately after, and at day 15 post-ischemia using laser doppler flowmetry of the plantar surface of the hindfoot. Relative perfusion (mean±SD) was calculated for in [M]Ctrl and [M]CD47 mice. One-way ANOVA.
